# Clonal lineage tracing and parallel multiomics profiling reveal transcriptional diversification induced by ARID1A deficiency

**DOI:** 10.1101/2025.05.24.655549

**Authors:** Kenichi Miyata, Liying Yang, Yuxiao Yang, Kohei Kumegawa, Reo Maruyama

**Affiliations:** Division of Cancer Epigenomics, Cancer Institute, Japanese Foundation for Cancer Research, Tokyo, Japan; Cancer Cell Communication Project, NEXT-Ganken Program, Japanese Foundation for Cancer Research, Tokyo, Japan; Cancer Cell Diversity Project, NEXT-Ganken Program, Japanese Foundation for Cancer Research, Tokyo, Japan

**Author notes:** **Correspondence to:** Reo Maruyama, M.D., Ph.D., Divison of Cancer Epigenomics, Cancer Institute, Japanese Foundation for Cancer Research 3-8-31, Ariake, Koto-ku, Tokyo, 135-8550, Japan, Kenichi Miyata, Ph.D., Divison of Cancer Epigenomics, Cancer Institute, Japanese Foundation for Cancer Research 3-8-31, Ariake, Koto-ku, Tokyo, 135-8550, Japan.

## Abstract

Phenotypic heterogeneity among genetically identical cancer cells underpins tumor progression and therapy resistance. However, the mechanism of epigenetic dysregulation that drives such divergence remains unclear. This study introduces integrated multimodal experimental platform specialized in analyzing inter/intraclonal heterogeneity (IMPACH), a scalable platform that integrates lineage tracing, genetic perturbation, and multimodal single-cell analysis. Utilizing IMPACH, results revealed that loss of epigenetic regulators—particularly ARID1A—enhances transcriptomic and epigenetic heterogeneity under controlled conditions. ARID1A deficiency promotes stochastic chromatin opening and induces an atypical gene expression program associated with poor clinical outcomes. Despite increased epigenetic randomness, chromatin changes remain localized to regulatory elements, partially linked to SMARCA4 binding sites. These findings revealed that chromatin remodeling defects promote clonal diversification through both stochastic and constrained mechanisms. Thus, owing to its scalability and versatility, IMPACH provides a robust framework for elucidating how epigenetic perturbations shape cellular heterogeneity in cancer.

## Introduction

Phenotypic heterogeneity—the nongenetic variation among genetically identical cancer cells—is critical in tumor progression and therapy resistance^1–3^. Individual cancer cells within clonal populations can adopt distinct transcriptional states, giving rise to subpopulations with varied proliferative capacities, metastatic potential, or drug sensitivity^4–6^. Although the origins of this heterogeneity are yet to be determined, one possible source is phenotypic plasticity—the ability of each cell to reversibly transition between functional states in response to intrinsic or extrinsic cues ^3, 7–10^. Strikingly, epigenetic modifiers that have emerged as key regulators of such plasticity dynamically reshape gene expression programs (GEPs) and chromatin landscapes^11^. Thus, we hypothesized that dysregulation of epigenetic modifiers may destabilize transcriptional control and enhance phenotypic divergence among genetically identical cells, driving cellular heterogeneity at the population level. However, directly dissecting how epigenetic dysregulation gives rise to such phenotypic heterogeneity is technically challenging.

Recent advances in single-cell technologies, such as single-cell RNA sequencing (scRNA-seq) and single-cell ATAC sequencing (scATAC-seq), have enabled high-resolution profiling of gene expression and chromatin states at the single-cell level ^12, 13^. Nonetheless, these approaches provide only static snapshots and cannot fully capture the dynamic processes underlying cell-state transitions. To delineate the contribution of epigenetic modifier dysregulation to phenotypic heterogeneity, transcriptomic and epigenomic changes at single-cell resolution must be monitored following targeted perturbations.

Hitherto, several methods that combine cellular barcoding with scRNA-seq or scATAC-seq to track clonal behavior alongside transcriptomic or chromatin state changes at the single-cell level have been developed^5, 6, 14–17^. Conversely, systems incorporating functional perturbations—such as CRISPR/Cas9 sgRNA or shRNAs—with single-cell omics analysis have enabled high-throughput characterization of how specific gene disruptions affect transcriptional or epigenomic states^18, 19, 20^, However, no existing platform has yet integrated clonal lineage tracing, targeted perturbation, and multimodal single-cell profiling into a cohesive framework (**Supplementary Table 1**).

To address this gap, integrated multimodal experimental platform specialized in analyzing inter/intra-clonal heterogeneity (IMPACH) was developed. This versatile platform combines single-cell-compatible barcoding with shRNA-based gene perturbation. IMPACH analyzes the influence of genetic perturbations, particularly of epigenetic regulators, on clonal composition, transcriptomic and chromatin states, and downstream functional consequences.

In this study, IMPACH was used, and the results demonstrated that knocking down epigenetic modifiers, such as ARID1A, generates biased clonal distributions characterized by a distinct MYC regulatory network. Notably, ARID1A knockdown induces dynamic chromatin remodeling and stochastic divergence in GEPs, giving rise to an ARID1A-dependent gene expression program, associated with poor clinical outcomes. Collectively, the findings highlight the utility of IMPACH as a robust platform for dissecting the mechanisms underlying epigenetic perturbation-mediated phenotypic heterogeneity in cancer.

## Results

### IMPACH reveals interclonal epigenetic and transcriptomic heterogeneity driven by the loss of specific epigenetic modifiers

Loss-of-function genetic aberrations in epigenetic modifiers are recurrently observed across various tumors^21, 22^. Given the essential role of these modifiers in maintaining proper epigenomic states^11^, we hypothesized that impaired epigenetic regulation, either through mutations or reduced expression, may contribute to increased phenotypic heterogeneity. Thus, TCGA pancreatic cancer RNA-seq data were analyzed, and transcriptomic diversity was quantified using two approaches—Shannon’s entropy^23^ and principal component analysis-based diversity indices^24^—revealing that tumors with reduced expression of epigenetic modifiers exhibited a high transcriptomic diversity (**Extended Data Figs. 1a–d**, see Methods for details). These results suggest that reduced expression of epigenetic modifiers is associated with high transcriptomic heterogeneity across tumors, implicating that functional impairment of these genes as a potential driver of phenotypic diversification in pancreatic cancer.

Furthermore, to assess this association and uncover the underlying mechanisms, we developed a lentiviral vector, shRNA-equipped pseudo-multiomics and cell tagging (shPseuMO-Tag) for lineage tracing, gene perturbation, and multimodal single-cell analysis via a DNA barcoding system (**Fig. 1a and Extended Data Figs. 2a, b**). Subsequently, an integrated analytical framework—IMPACH was established to analyze phenotypic heterogeneity in a flexible and scalable manner. IMPACH offers adjustable experimental configurations by tuning the clone number, culturing strategy (pooled or isolated), and expansion time, enabling flexible analysis from clonal dynamics to inter-/intraclonal variability (**Extended Data Fig. 3**; see Methods for details). As a proof of concept, IMPACH was used to investigate how the reduced function of representative epigenetic modifiers affects transcriptomic and epigenomic heterogeneity. Four modifiers—KDM6A, SETD2, ARID1A, and SMARCA4—were selected based on their recurrent loss-of-function mutations in pancreatic cancer and established links to poor clinical outcomes^25–30^, as well as their association with increased transcriptomic diversity in tumors with low expression levels (**Extended Data Figs. 1c, d**).

**Fig. 1.**
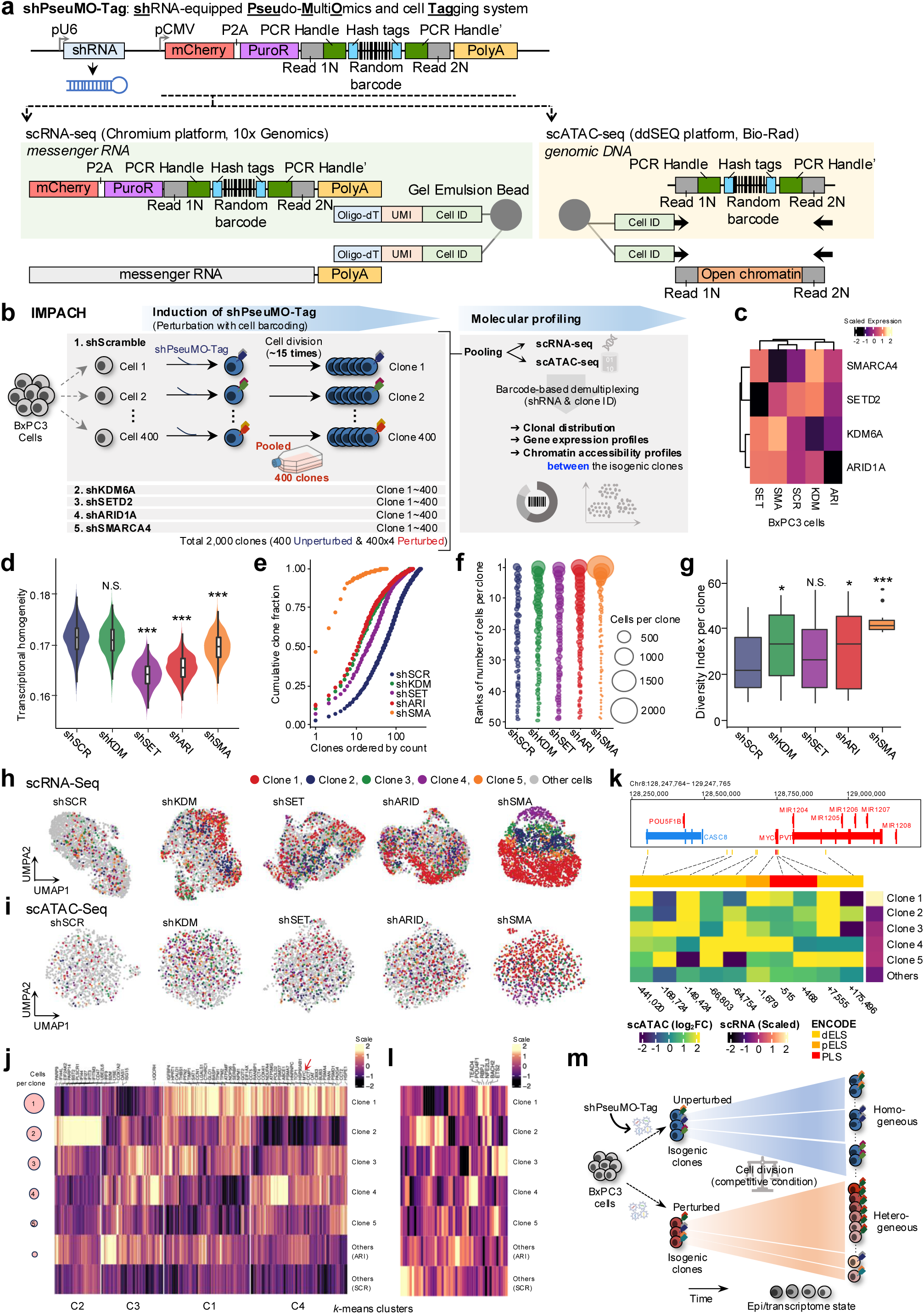
Development of the shPseuMO-Tag and multiomics analysis reveals clone-specific epigenetic and transcriptomic states. **a**, Schematic of the shPseuMO-Tag (shRNA-equipped pseudo-multiomics and cell tagging) vector, which combines an shRNA module with a multiomics-compatible DNA barcode embedded in the 3′ UTR of an mCherry–PuroR fusion gene. **b**, Experimental workflow. BxPC3 cells transduced with shPseuMO-Tag targeting KDM6A, SETD2, ARID1A and SMARCA4, or scramble control, were clonally expanded (approximately 15 divisions), pooled, and subjected to scRNA-seq and scATAC-seq using the same cellular mixture. **c**, Expression levels of genes targeted by each shPseuMO-Tag. **d,** Transcriptomic heterogeneity assessed using normalized mutual information (NMI). For each perturbation, 1,000 bootstrap samples of 50 cells were analyzed. ***P < 0.001 by one-way ANOVA followed by Dunnett’s post hoc test. **e**, Cumulative distribution of barcode frequencies ranked by abundance. **f**, Proportional bubble plot represents the top 50 most abundant clones. **g**, Transcriptomic heterogeneity analyzed using principal component analysis (PCA) on clone-level profiles. ***P < 0.001, *P < 0.05; NS, not significant by one-way ANOVA followed by Dunnett’s post-hoc test. **h, i**, UMAP of both scRNA-seq (**h**) and scATAC-seq (**i**) cells with cells from the top five abundant clones highlighted. **j**, Heatmap of 525 differentially expressed genes across individual clones compared with SCR controls, showing scaled expression values. **k**, Heatmap of chromatin accessibility (log₂ fold-change to SCR controls) in the top five dominant clones and remaining minor clones. The top annotation indicates ENCODE cCRE classification; right panel shows MYC expression levels. **l**, Heatmap of transcription factor expression profiles predicted to bind MYC-associated cis-regulatory elements. **m**, ARID1A loss disrupted clonal balance, promoting the emergence of dominant clones with distinct transcriptomic and chromatin accessibility states.

Herein, the human pancreatic cancer cell line BxPC3, was transduced with shPseuMO-Tag constructs targeting each gene, along with a nontargeting scramble control (**Fig. 1b and Extended Data Fig. 4a**). Twenty-four hours post-transduction, 400 cells/dish (approximately 400 clones) were seeded and cultured for approximately 15 cell divisions. The expanded populations were pooled and subjected to parallel scRNA-seq and scATAC-seq to profile approximately 2,000 clones (400 clones × five shRNAs, including control) simultaneously with minimal batch effects^31^ (**Fig. 1b**). Following demultiplexing based on shRNA identifier sequences, 2500–3500 cells per shRNA condition were obtained in the scRNA-seq dataset and 1500–2500 cells per shRNA condition in the scATAC-seq dataset (**Extended Data Figs. 4b, c**). Knockdown of each target gene was confirmed by scRNA-seq, validating both the successful perturbation induced by shPseuMO-Tag system and the accuracy of the PseuMO-Tag Decoder pipeline (**Fig. 1c and Extended Data Fig. 2c**; see Methods for details).

To assess how reduced epigenetic modifier function affects heterogeneity, normalized mutual-information^5, 32^ and entropy-based diversity indices^33^ were calculated in mixed clonal populations (see Methods). Knockdown of SETD2, ARID1A, and SMARCA4 increased both transcriptomic and epigenomic heterogeneity levels compared with the control (**Fig. 1d and Extended Data Fig. 4d**). Subsequently, individual clones were demultiplexed using the PseuMO-Tag barcode (**Extended Data Figs. 4b,c,e–h**). Although 415 clones were detected in the control condition, the number of identifiable clones was substantially lower in the knockdown conditions (KDM6A, 249; SETD2, 272; ARID1A, 262; SMARCA4, 60), indicating clonal imbalance and selective expansion of specific clones (**Figs. 1e–f and Extended Data Fig. 4i**). Notably, these shifts emerged under standard culture conditions without additional external selection pressures, suggesting that knocking down epigenetic modifiers enhances intrinsic phenotypic and functional heterogeneity, thereby leading to differential clonal fitness.

Furthermore, to assess the utility of IMPACH, the transcriptomic and epigenomic profiles of dominant vs. minor clones in each knockdown condition were compared, and substantial interclonal variations were observed (**Figs. 1g–i**). In the ARID1A-knockdown group, the most dominant clone exhibited high expression of MYC and its downstream targets and epithelial–mesenchymal transition (EMT)-related genes (*C4* and *C1*; **Fig. 1j**), whereas other clones were enriched with interferon (IFN)- or oxidative phosphorylation-related genes (*C2* or *C3*; **Fig. 1j and Extended Data Figs. 5a–d**). MYC and EMT pathways are associated with aggressive features in pancreatic cancer^34, 35^; previous studies have linked ARID1A loss to MYC upregulation^28^ and EMT activation^36^.

Subsequently, to explore regulatory changes associated with ARID1A loss, chromatin accessibility around the MYC locus was examined. scATAC-seq revealed increased accessibility, in a dominant clone-specific manner, at a candidate cis-regulatory element (cCRE) approximately 441 kbp upstream of the MYC transcription start site, within the same topologically associating domain (TAD) (**Fig. 1k and Extended Data Fig. 5e**). Motif analysis predicted the potential binding sites for ETS2 and TBP at this region, and both factors were upregulated in the dominant clone (**Fig. 1l and Extended Data Fig. 5f**). Given that MYC is a known transcriptional target of ETS2^37, 38^, these findings suggest that ARID1A loss promotes clonal dominance through ETS2-mediated MYC transcriptional activation.

Together, these results demonstrated that IMPACH enables the simultaneous analysis of clonal composition dynamics, transcriptomic states, and chromatin accessibility at the clonal level (**Fig. 1m**). Moreover, this high-resolution platform allows parallel evaluation of perturbation effects across hundreds of clones while minimizing batch effects^31^.

### IMPACH identifies stochastic divergence in transcriptional states driven by ARID1A loss

Interestingly, ARID1A showed a pronounced increase in clonal diversity (**Fig. 1g and Extended Data Fig. 5c**) and is frequently mutated in pancreatic^39^ and other cancers^40^, indicating that this dysfunctional epigenetic modifier contributes to intraclonal phenotypic heterogeneity. Although the initial experiment involved approximately 2000 clones (**Fig. 1b**), the limited number of cells per clone in single-cell analyses hindered a detailed assessment of intraclonal variations. Therefore, in the next phase, we reduced the number of clones to fewer than 30 and established them through single-cell cloning to ensure separation and minimize competition. This setup enabled us to observe phenotypic variation within individual clones under minimal extrinsic selection pressure and allowed us to track or manipulate specific clones in downstream functional assays (**Fig. 2a**).

**Fig. 2.**
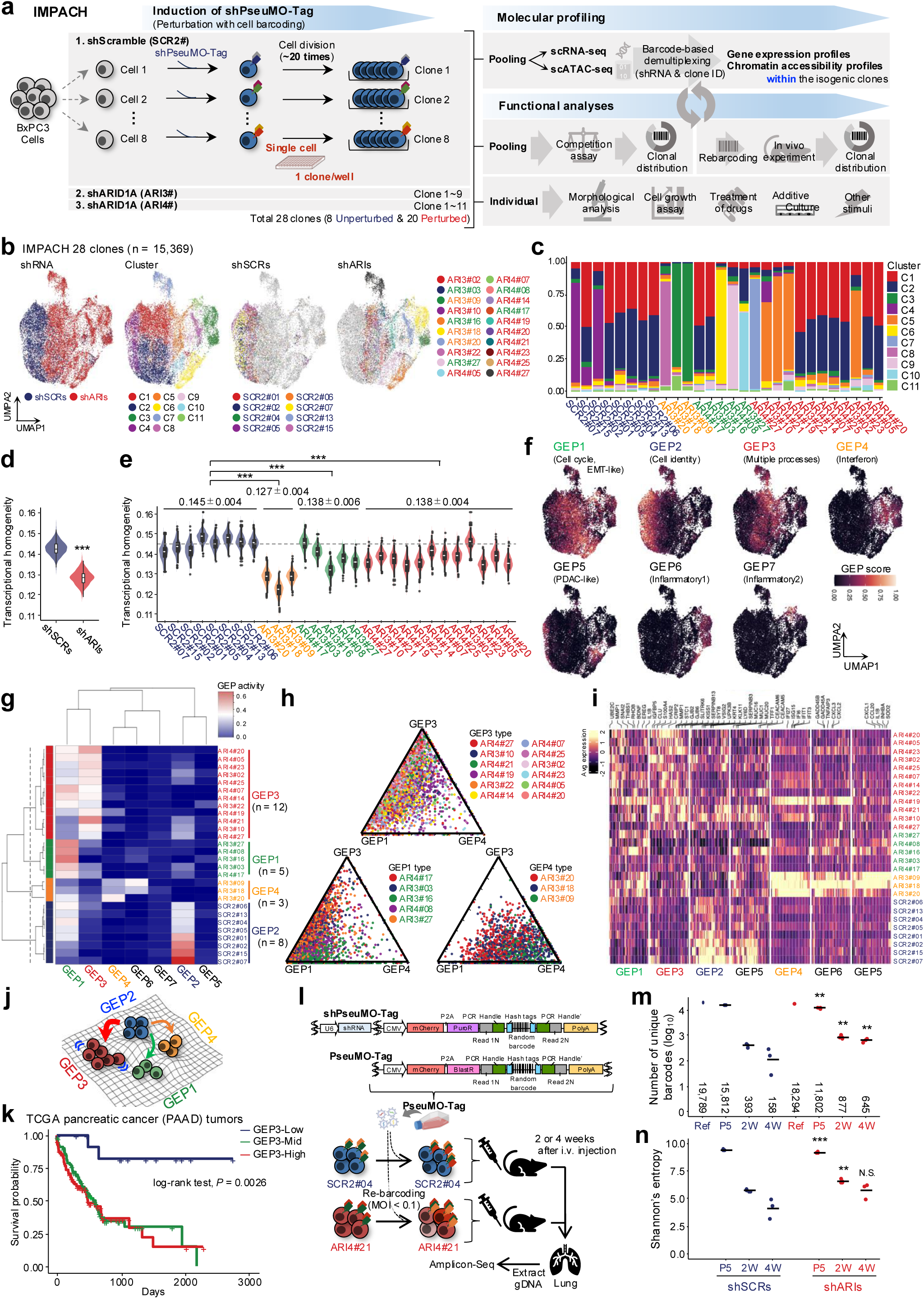
IMPACH reveals clone-specific transcriptional programs and stochastic divergence driven by ARID1A loss. **a**, Schematic of the IMPACH system. BxPC3 cells were independently transduced with shScramble or shARID1A constructs, cloned, and expanded (approximately 20 divisions). Single-cell transcriptomic and epigenomic profiling, as well as functional assays, were performed on individual clones. **b**, UMAP projections of 15,369 single cells derived from 28 clones, colored by shRNA type, transcriptomic clusters, or individual clone identity for shScramble and shARID1A conditions. **c**, Distribution of cells across shRNA types (**c**) and clonal compositions across transcriptomic clusters (**d**). **d**, **e**, Transcriptomic heterogeneity quantified by normalized mutual information (NMI), calculated per shRNA type (**d**) or individual clone (**e**). Values represent mean ± standard deviation according to gene expression program (GEP) type assignment; dashed line indicates the median of control clones. **f**, UMAP visualizations of GEP activity scores. **g**, Hierarchical clustering of GEP activity scores across clones, with values representing the median activity per clone. **h**, Ternary plots showing the distribution of ARID1A-knockdown clones across GEP1–GEP3. Each dot is a cell colored according to the GEP type assignment. **i**, Heatmap showing the average expression per clone of the top 50 genes defining each GEP. **j**, Conceptual model illustrating stochastic transcriptional divergence among isogenic clones driven by ARID1A loss. **k**, Kaplan–Meier analysis of the overall survival in the TCGA PAAD cohort, stratified by the average expression of GEP3-associated genes (low, mid, high). *P*-value calculated using the log-rank test. **l**, Schematic of PseuMO-Tag-compatible re-barcoding strategy. SCR2#04 and ARI4#21 clones were labeled with PseuMO-Tag barcodes and injected intravenously into NSG mice. **m**, **n**, Number of unique barcodes (**m**), and Shannon’s entropy of barcode distributions (**n**) detected in lung tissues or in vitro cultures at 2 and 4 weeks after injection. Each dot represents a biological replicate; bars indicate the median. Comparison between matched clones (SCR2#04 vs ARI4#21). **P < 0.01, ***P < 0.001 by Wilcoxon rank-sum test.

shPseuMO-Tag constructs targeting ARID1A (two independent shRNAs: ARI3# and ARI4#) or a nontargeting control (SCR2#) were introduced into BxPC3 cells. Following single-cell cloning and approximately 20 cell divisions, 28 clonal subpopulations were established (8 controls and 20 ARID1A knockdowns; **Extended Data Fig. 6a**). Morphological and proliferative analyses showed that ARID1A-dificient clones displayed broader morphological diversity (types 3–6) and more variable, generally reduced growth compared with controls (**Extended Data Figs. 6b, c**), suggesting that ARID1A-deficiency enhances phenotypic heterogeneity among independently derived clones, thus reflecting pre-existing variability within the parental BxPC3 population.

Subsequently, all 28 clones were pooled into a single cellular mixture, and scRNA-seq and scATAC-seq were performed to minimize the batch effect^31^ while evaluating transcriptional variations within and across clones (**Fig. 2a**). Clone identities were assigned via PseuMO-Tag barcodes (**Extended Data Figs. 2c and 6d**). scRNA-seq revealed distinct transcriptional shifts in ARID1A-deficient clones compared with controls (**Figs. 2b, c and Extended Data Figs. 6e, f**). Notably, ARID1A-deficient clones not only exhibited higher overall transcriptomic diversity (**Fig. 2d**), but also showed substantial variation in the transcriptional diversity between individual clones (**Fig. 2e**).

Then, consensus non-negative matrix factorization (cNMF)^41^ was applied to the scRNA-seq data to further dissect ARID1A loss-associated transcriptional changes, and seven GEPs were identified (**Figs. 2f, g, Extended Data Figs. 7a, b**, and **Supplementary Table 2**). In control clones, GEP2 reflected BxPC3-specific cellular identity (**Extended Data Fig. 7c**), whereas ARID1A-deficient clones showed enrichment for multiple distinct programs, including an EMT-like (GEP1) and an IFN-related (GEP4) programs (**Figs. 2g–i and Extended Data Fig. 7d**). Based on GEP compositions, ARID1A-deficient clones were segregated into three groups: GEP4/6-enriched (yellow), GEP1-enriched (green), and GEP3-dominant (red) clusters (**Figs. 2g–i**). These findings are consistent with the results of previous studies linking ARID1A loss to EMT activation^36, 42, 43^ and inflammatory signaling^44, 45^, and suggest that ARID1A deficiency induces stochastic, clone-specific divergence in GEPs at the single-cell level (**Fig. 2j**).

Among the GEPs identified by cNMF, GEP3 has emerged as a unique program in ARID1A-deficient clones. It did not strongly associate with any single hallmark pathway but showed weak enrichment across multiple processes, including coagulation, gap junction assembly, and hyaluronan biosynthetic processes (**Figs. 2g–j and Extended Data Figs. 7d, e**). GEP3 included genes such as CLU, a regenerative marker of pancreatic and intestinal epithelia^46^, and gap junction components (GJB2, GJB6). Notably, GEP3 activity was stable across at least 10 cell divisions (**Extended Data Fig. 7f**), indicating the heritable nature of ARID1A-induced GEPs.

Further, to test the functional relevance of GEP3, a lung colonization assay was performed by co-injecting five control and five ARID1A-deficient clones (**Extended Data Figs. 8a, b**). Competition between the clones was assessed based on their relative engraftment capacities. ARID1A-deficient clones exhibited significant engraftment capacity. Hence, to exclude the possibility that rare pre-adapted subclones accounted for this advantage, a refined re-barcoding system (PseuMO-Tag) with modified selection markers was developed (**Fig. 2l**), and one control (SCR2#04) and one GEP3-high clone (ARI4#21) were re-barcoded (**Fig. 2m, Extended Data Figs. 8c, d**). The barcode complexity remained largely stable after five in vitro passages (**Extended Data Fig. 8e**). However, in the lung metastasis model, a strong selective outgrowth was observed (**Extended Data Figs. 8f, g**). Notably, ARI4#21 exhibited strong clonal outgrowth, outperforming SCR2#04 in the number of successfully engrafted subclones and exhibited higher Shannon’s entropy (**Fig. 2n**). These findings implied that GEP3-high clones possess a selective advantage in ectopic colonization.

Finally, TCGA pancreatic cancer RNA-seq data were analyzed to evaluate the potential clinical relevance of GEP3. Tumors with high GEP3-related gene expression are associated with poorer patient survival (**Fig. 2k**), suggesting that GEP3 may contribute to aggressive tumor phenotypes. Collectively, these findings highlight the potential clinical and functional significance of ARID1A-driven transcriptional divergence.

### ARID1A knockdown increases the diversity of chromatin accessibility

Since ARID1A deficiency induced divergent GEPs, the associated changes in chromatin accessibility were investigated. Using the same mixture pooled for scRNA-seq (**Fig. 2a**), scATAC-seq was performed, and clonal identities were determined via PseuMO-Tag decoding (**Extended Data Fig. 9a**). UMAP analysis revealed 10 distinct chromatin clusters (C1–C10), with ARID1A-deficient clones broadly distributed across clusters, whereas scramble clones remained confined (**Figs. 3a–c, Extended Data Figs. 9b–d**), suggesting chromatin landscape remodeling upon ARID1A loss. Clone-level multiomics analysis integrating scATAC-seq and scRNA-seq revealed that clones enriched for specific GEPs aligned with distinct chromatin clusters. IFN-high clones (yellow, GEP4/6) predominantly occupied C2, EMT-high clones (green, GEP1) were distributed across C1, C2, C5, and C10, and GEP3-associated clones (red) mainly clustered into C3 and C4. Notably, C3 and C4 exhibited numerous accessible peaks (**Extended Data Fig. 9c**), consistent with the atypical and functionally uncharacterized transcriptional signature of GEP3 clones.

**Fig. 3.**
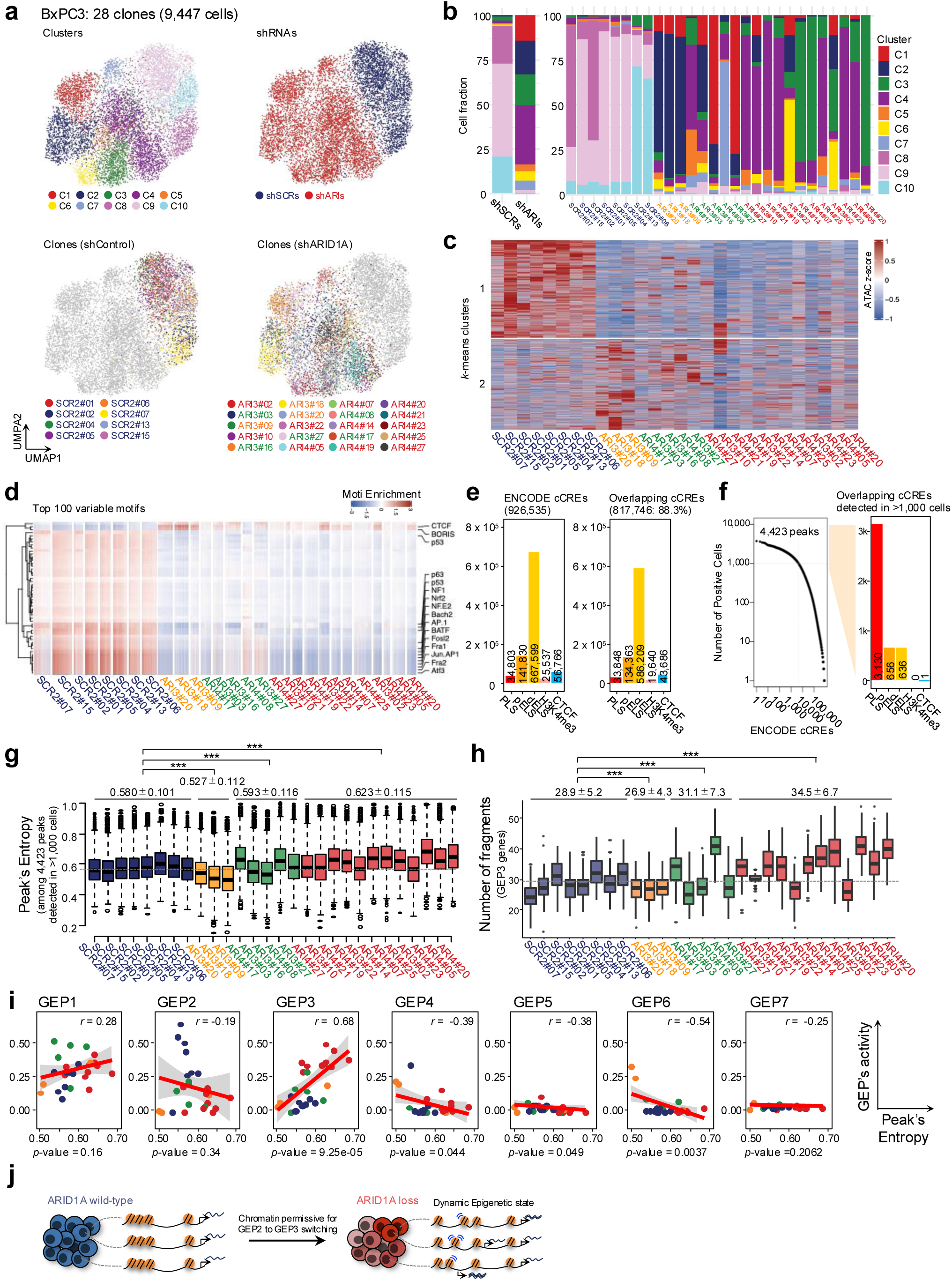
ARID1A loss alters chromatin accessibility and increases epigenetic diversity. **a**, UMAP projections of scATAC-seq profiles from 9,447 cells (28 clones), colored by unsupervised chromatin clusters (top left), shRNA type (top right), and clone identity for shScramble (bottom left) and shARID1A (bottom right) conditions. **b**, Distribution of cells across shRNA types (left) and clonal compositions across chromatin clusters (right). **c**, Heatmap of chromatin accessibility (z-score) for 7,202 differentially accessible regions (FDR ≤ 0.5, log₂FC ≥ 0.25) across clones, grouped by k-means clustering. **d**, Motif enrichment analysis of the top 100 most variable transcription factor motifs across clones. **e**, Overlap between identified ATAC peaks and ENCODE candidate cis-regulatory elements (cCREs; total = 926,535). Of these, 88.3% overlapped with at least one ATAC peak. **f**, Number of cells with positive chromatin accessibility at ENCODE cCREs (left) and the number of overlapping cCREs detected in >1,000 cells (right). **g**, Peak entropy scores calculated per clone across 4,423 shared peaks. Clones are colored by GEP cluster identity. Values represent mean ± standard deviation according to GEP type assignment; dashed line indicates the median of control clones. ***P < 0.001 by one-way ANOVA followed by Dunnett’s post-hoc test. **h**, Number of ATAC fragments mapped to the regulatory regions of GEP3-related genes per clone. Each dot represents one of 100 bootstrap replicates; dashed line indicates the median of control clones. ***P < 0.001 by one-way ANOVA followed by Dunnett’s post-hoc test. **i**, Correlation between peak entropy and GEP activity (GEP1–GEP7) across clones are represented by Pearson’s correlation coefficient (*r*) and associated *P*–values. **j**, Conceptual model. ARID1A loss destabilizes chromatin accessibility and increases epigenetic potential, enabling stochastic shifts from GEP2 (BxPC3 cell identity) to GEP3 (multiple processes).

Intriguingly, ARID1A-deficient clones, particularly those associated with high GEP3 activity (red group), exhibited a global reduction in the transcription factor motif enrichment scores (**Fig. 3d and Extended Data Figs. 9e, f**). This widespread decrease suggests that ARID1A deficiency erodes pre-existing chromatin regulatory networks, leading to increased stochastic and less structured patterns of chromatin accessibility. Despite increased stochasticity, 88.3% of the accessible regions overlapped with ENCODE cCREs (**Fig. 3e**), indicating that chromatin opening remained largely confined to functional loci. Peak entropy analysis using 2,234 ENCODE cCREs confirmed greater variability in chromatin accessibility in ARID1A-deficient clones than in controls; GEP3-associated clones showed the highest entropy (**Figs. 3f, g**). Multiomics analysis confirmed increased accessibility of GEP3-high clones at the promoter sites of GEP3-related genes (**Fig. 3h**). Moreover, GEP3 activity was strongly correlated with peak entropy (**Fig. 3i**; *r* = 0.68).

Collectively, these results suggest that ARID1A deficiency destabilizes chromatin accessibility, promoting stochastic yet functionally constrained epigenomic shifts that facilitate GEP3-related transcriptional activation (**Fig. 3j**).

### Association of SMARCA4-bound chromatin states with GEP3 activity under ARID1A-deficient conditions

Furthermore, to characterize the regulatory landscape associated with GEP3 activation, we leveraged scRNA-seq data to infer transcriptional regulatory activities across clones (**Fig. 4a**). Unsupervised clustering stratified the clones into three major groups: one dominated by IRF-family transcription factors associated with IFN-high clones, another resembling shScramble controls, and the third encompassing clones enriched for GEP3 activity. Notably, the SMARCA4 activity was increases across ARID1A-deficient clones, including both GEP3-high and IFN-high populations, compared with controls. This observation suggested that ARID1A loss enhances SMARCA4-associated regulatory features, potentially contributing to the altered chromatin landscape underlying divergent transcriptionalprograms such as GEP3.

**Fig. 4.**
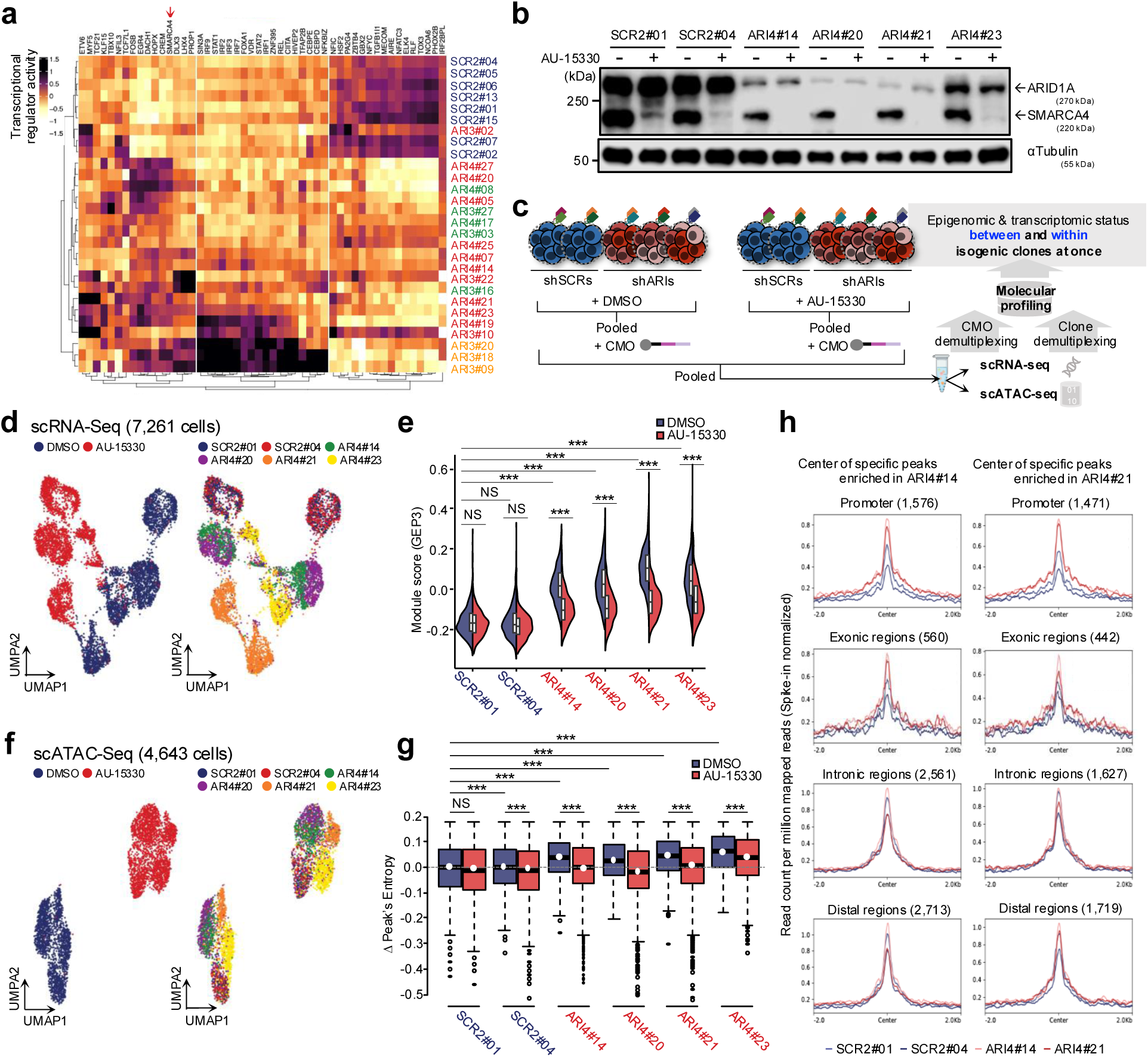
SMARCA4 partially modulates GEP3 activity and epigenetic diversity under ARID1A-deficient conditions. **a**, Heatmap showing transcriptional regulatory activity scores across clones. The top 50 genes ranked by variability are displayed. **b**, Immunoblot analysis of ARID1A, SMARCA4, and α-tubulin in selected clones treated with either DMSO or the SMARCA4 inhibitor AU-15330. **c**, Experimental workflow for single-cell multiomics analysis under SMARCA4 inhibition. shScramble and ARID1A-knockdown clones were treated with either DMSO or AU-15330, pooled, and barcoded using cell multiplexing oligos (CMOs), followed by scRNA-seq and scATAC-seq. **d**, UMAP plots of 7,261 cells profiled by scRNA-seq, and colored by treatment condition (left) or clone identity (right). **e**, Violin plots of GEP3 activity across individual clones treated with DMSO or AU-15330. ***P < 0.001, **P < 0.01, *P < 0.05; NS, not significant by one-way ANOVA followed by Tukey’s post-hoc test. **f**, UMAP plots of 4,643 cells profiled by scATAC-seq and colored by treatment condition (left) or clone identity (right). **g**, Box plots of peak entropy scores in clones treated with DMSO or AU-15330. ***P < 0.001, **P < 0.01, *P < 0.05; NS, not significant by one-way ANOVA followed by Tukey’s post-hoc test. **h**, CUT&Tag signal intensity profiles for peaks specifically enriched in ARI4#14 (left) and ARI4#21 (right). Peaks were categorized into promoter, exonic, intronic, and distal regions. Lines represent average read counts per million mapped reads, normalized by spike-in controls.

The functional significance of these SMARCA4-associated regulatory features in ARID1A-deficient was explored via the protein depletion experiments using AU-15330 in selected clones (controls, #1 and #4; GEP3-high clones, #14, #20, #21, and #23) (**Fig. 4b**). The clones were evaluated using multiomics on clonal pools treated with DMSO or AU-15330; the individual clones were distinguished by cell multiplexing oligos and utilizing the advantages of shPseuMO-Tag (**Fig. 4c**). scRNA-seq showed that high GEP3 activity in the four GEP3-high clones was attenuated following SMARCA4 depletion (**Figs. 4d, e and Extended Data Figs. 10a–d**). Consistently, scATAC-seq revealed that the high peak entropy in GEP3-high clones decreased upon SMARCA4 depletion (**Figs. 4f, g and Extended Data Figs. 10e–h**). These results suggest that SMARCA4 maintains the features of the GEP3-associated state under ARID1A-deficient conditions.

To further investigate the correlation between SMARCA4 binding and GEP3-related transcriptional changes, CUT&Tag was performed to map the genome-wide SMARCA4 binding sites. SMARCA4 binding landscapes were markedly altered in ARID1A-deficient clones (**Extended Data Figs. 10i, j**). In GEP3-high clones, SMARCA4 binding was enriched at promoters and exonic regions of GEP3-associated genes (**Fig. 4h**). These observations suggest that chromatin regions associated with SMARCA4 binding become more accessible following ARID1A deficiency, potentially facilitating the activation of GEP3-related transcriptional programs.

## Discussion

This study introduced IMPACH, a scalable multimodal platform for dissecting how epigenetic perturbations influence clonal and intraclonal heterogeneity in cancer. Thus, integrating clonal barcoding, gene perturbation, and single-cell multi-omics profiling within a unified framework enables high-resolution analyses of clonal behavior, transcriptomic plasticity, and chromatin remodeling under controlled genetic backgrounds.

A key strength of IMPACH is its flexible scalability, which was demonstrated through two complementary proof-of-concept experiments. First, approximately 2,000 clones were profiled simultaneously, and results revealed how ARID1A knockdown induces clonal biases characterized by distinct transcriptomic and epigenomic features. Second, the number of clones was reduced to <30 via single-cell cloning, allowing an in-depth analysis of intraclonal heterogeneity. Importantly, in both settings, each clone represents a lineage derived from an individual perturbation event at the time of shRNA introduction, enabling precise tracking of perturbation-induced phenotypic diversification. Moreover, IMPACH’s pooled single-cell analysis minimized batch effects, reduced experimental costs and labor, and allowed direct comparison of numerous clones within a single experiment. In addition, barcoding allowed the integration of transcriptomic and epigenomic profiles at clonal resolution, leading to the concept of “pseudo-multiomics”.

In the background of ARID1A deficiency, loss of chromatin remodeling activity destabilized pre-existing regulatory networks, increased stochastic chromatin opening, and drove the emergence of divergent transcriptional programs, including GEP3 associated with metastatic potential. Although chromatin remodeling appeared random, the accessible regions remained largely constrained to functional cis-regulatory elements. Further analysis revealed that SMARCA4, another SWI/SNF component, was enriched at GEP3-associated loci, and its depletion partially attenuated GEP3 activity. These findings indicate that, despite ARID1A loss, residual chromatin regulatory activities—at least partially mediated by SMARCA4—may bias stochastic chromatin opening toward biologically meaningful regions. Nonetheless, the precise mechanisms underlying this interplay between different chromatin remodelers warrant further investigation.

In conclusion, IMPACH offers a robust and scalable framework to link epigenetic perturbations with cellular heterogeneity, providing insights into how dysregulated chromatin remodeling promotes phenotypic plasticity and clonal diversification in cancer. Its versatility also makes it applicable to diverse contexts, including treatment resistance, metastasis, and lineage plasticity across cancer types.

## Methods

### Cell culture and reagents

BxPC3 cells were cultured in RPMI-1640 medium (Nacalai Tesque) supplemented with 10% fetal bovine serum and penicillin/streptomycin (Sigma-Aldrich) in a 5% CO_2_ incubator at 37°C. All cell lines used tested negative for mycoplasma. Some BxPC3 clones were incubated with 1 μM AU-15330 (Selleck) or DMSO (Sigma-Aldrich) for 24 h (western blot and scRNA-seq) or 4 h (scATAC-seq).

### Cell proliferation

To measure cell proliferation, BxPC3 clonal populations were seeded in triplicate at a density of 1,000 cells per well in 96-well plates (Corning). The cells were cultured for 7 days, during which phase-contrast images were acquired every 24 h using a 4× objective on an IncuCyte SX5 Live-Cell Analysis System (Sartorius). Furthermore, confluence was automatically quantified at each time point using the integrated IncuCyte analysis software.

### Morphological classification of the clonal populations

To assess the morphological heterogeneity among the clonal populations, bright-field images were captured for each clone using a BZ-X700 microscope (Keyence). The clones were classified into six predefined morphological types (Types 1–6) based on a stepwise decision-making process. First, if the average cell diameter was ≥20 μm, the clone was assigned to Type 6. If this criterion was not met, colony-level features were assessed. Clones, in which cells formed tightly packed and condensed colonies, were categorized as Type 2. Clones with looser intercellular contacts and scattered arrangements, which were suggestive of a mesenchymal-like morphology, were classified as Type 5. If the cells appeared flattened, with an enlarged surface area, prominent perinuclear halos, or surface wrinkling, the clone was assigned to Type 4. Clones containing cells with an average diameter exceeding 11 μm—larger than that of the parental line—were designated as Type 3. Clones that did not meet these criteria and resembled the morphology of the parental cell line, characterized by cohesive growth and an epithelial-like appearance, were categorized as Type 1.

### RT-qPCR

Total RNA was extracted using the RNeasy Plus Mini Kit (Qiagen) based on the manufacturer’s instructions. Reverse transcription was done using the PrimeScript RT Master Mix (TaKaRa). Quantitative PCR was carried out on a StepOne Plus Real-Time PCR System (Thermo Fisher Scientific) using SYBR Premix Ex Taq II (Tli RNaseH Plus; TaKaRa, #RR820A), as previously described^47–49^. The following primer sequences were used: KDM6A: 5′-GCGGACAAAAGAAGAACCAG-3′ (forward) and 5′-AGATGAGGCGGATGGTAATG-3′ (reverse); SETD2: 5′-AAACACAACAACTGAACGAGG-3′ (forward) and 5′-GAGTTTGCTTGTCTGGGTCTC-3′ (reverse); ARID1A: 5′-CGGCCAATGGATGGCACATA-3′ (forward) and 5′-TCGGCCAAACTGGAATGGAA-3′ (reverse); SMARCA4: 5′-CTCGGTCCGTCAAAGTGAAGA-3′ (forward) and 5′-AGCGGTCCTCCTCTTGTTCC-3′ (reverse); and ACTB (housekeeping gene): 5′-AGAGCTACGAGCTGCCTGAC-3′ (forward) and 5′-AGCACTGTGTTGGCGTACAG-3′ (reverse). Relative gene expression was calculated using the ΔΔCt method and normalized to ACTB expression.

### Western blot analysis

Western blot analysis was performed as previously described^47, 48, 50^ with minor modifications. Cell pellets were lysed in buffer containing 0.1 M Tris-HCl (pH 7.5), 10% glycerol, and 1% SDS, boiled for 5 min, and centrifuged at 15,000 rpm for 10 min. Protein concentrations were determined using the BCA Protein Assay Kit (Pierce). Equal amounts of protein were separated by SDS–PAGE and transferred onto PVDF membranes (Millipore). The membranes were blocked with 5% skim milk (Megumilk) or 5% bovine serum albumin (Sigma-Aldrich) in tris-buffered saline containing 0.1% Tween 20 (TBST). They were incubated overnight at 4°C with the following primary antibodies diluted in blocking buffer: anti-ARID1A (Cell Signaling Technology, #12354, 1:1,000), anti-BRG1 (Cell Signaling Technology, #49360, 1:1,000), and anti-α-tubulin (Sigma-Aldrich, Clone B-5-1-2, T5168, 1:5,000). After washing three times in TBST, the membranes were incubated with HRP-conjugated secondary antibodies, which included either anti-mouse IgG (Cell Signaling Technology, #7076, 1:5,000) or anti-rabbit IgG (Cell Signaling Technology, #7074, 1:5,000), for 1 h. Moreover, signals were visualized using the SuperSignal West Femto Maximum Sensitivity Substrate (Thermo Fisher Scientific) and captured with an Odyssey Fc Imaging System (LI-COR).

### Plasmid construct

The shPseuMO-Tag vector contains a DNA barcode in the 3′ UTR of a polyA-tagged mCherry–PuroR fusion construct, flanked by Tn5 adaptor sequences. This design enables simultaneous capture of the same barcode during 3′ Chromium scRNA-seq (10x Genomics) and ddSEQ scATAC-seq (Bio-Rad) without requiring custom reagents or protocols.

The shPseuMO-Tag plasmid was constructed based on the pLenti CMV GFP Puro (658-5) vector (Addgene, #17448). The cPPT/CTS, CMV enhancer, CMV promoter, and EGFP regions were removed using *HpaI* and *SalI* restriction enzymes. Next, the U6 promoter, shRNA sequence (flanked by *BamHI* and *SalI* restriction sites), cPPT/CTS, CMV enhancer, CMV promoter, and a mCherry-P2A-PuroR-Read N1-PCR handle1-Stuffer (for inserting cell barcode sequences)-PCR handle2-Read N2-SV40 poly(A) signal cassette were inserted via In-Fusion 5X HD Cloning Plus (Takara Bio Cat #638909), using 20– 30 bp homologous sequences, which resulted in shPseuMO-Tag-Stuffer (available upon request). The full-length sequence is provided in Supplementary Table 3.

The sequences used for shRNA were as follows: KDM6A: 5′-GATGCAAGTC TATGACCAAT T-3′ (Target sequence of pre-designed shRNA) (TRCN0000107763, Sigma-Aldrich); SETD2: 5′-AAACCAACAG TCTGTCAGTG TA-3′^51^; ARID1A: 5′-GCCTGATCTA TCTGGTTCAA T-3′ (ARI3) and 5′-GCATCCTTCC ATGAACCAAT C-3′ (ARI4)^52^; and SMARCA4: 5′-CGGCAGACAC TGTGATCATT T-3′^53^. Nontargeting shRNA (scramble) sequence was used as negative control: 5′-AGTCTTAATC GCGTATAAG-3′^52^. The hairpin sequence used was as follows: 5′-CTGTGAAGCC ACAGATGGG-3′^54^. To assemble the cell barcode, based on a slight modification of a previous report^16^, a 59-bp oligonucleotide containing a 10-bp random sequence flanked at both ends by a 5-bp HashTag sequence and a reverse extension primer was obtained from Integrated DNA Technologies: 5′-GAGCCTCGTC TCCTGACTNN NNNNNNNNNN NNNNNNNNCG TTGTGAGACG CATGCTGCA-3′. The 5-bp HashTag sequences were as follows: shSCR: ATGCA; shKDM: AGCTG; shSET: TGCTA; shARID: CTGAA; shSMA: GCCAA; shSCR2: ATGCA; shARID3: CGTGC; shARID4: CTGAA. The following extension reaction was performed to generate double-stranded barcode-sgRNA oligonucleotides: 25 µl NEBNext® High-Fidelity 2X PCR Master Mix (NEB, M0541), 1 µl of 100 µM cell barcode, 2 µl of 100 µM reverse extension primer (5′-TGCAGCATGC GTCTCACAAC G-3’), and 22 µl water. The reaction conditions were as follows: 98°C for 2 min, 10× (65°C for 30 s, 72°C for 10 s), 72°C for 2 min, and hold at 4°C. The double-stranded barcode-sgRNA oligonucleotide was purified using a QIAquick PCR Purification kit (QIAGEN, 28104). The double-stranded product contained two *BsmBI* sites that, upon digestion, generated complementary overhangs for ligation into shPseuMO-Tag-Staffer. Next, 1.5 µg of BsmBI-v2 digested (New England Biolabs, R0739) shPseuMO-Tag-Staffer vector was added to a Golden Gate Assembly reaction with the double-stranded barcode insert at a molar ratio of 1:5 and cycled 100× (42°C for 2 min and 16°C for 5 min). The reaction was purified and concentrated in 10 µl of water using the DNA Clean & Concentrator kit (Zymo, D4033) and transformed into NEB® Stable Competent *E. coli* (New England Biolabs, C3040). Transformants were inoculated into 500 mL of LB medium containing 100 µg mL^−1^ ampicillin and incubated overnight at 30°C. Bacterial cells were collected by centrifugation at 6,000 RCF at 4°C for 15 min, and plasmid DNA was extracted using a QIAGEN Plasmid Plus Maxi kit (QIAGEN, 12943).

The PseuMO-Tag vector was constructed by removing the shRNA cassette from the shPseuMO-Tag via *HpaI* and *SalI* digestion, followed by a blunting reaction. The PuroR sequence was replaced with a BlastR sequence cloned from lentiCRISPR v2-Blast (Addgene, #83480) using In-Fusion cloning. For barcode construction, a 77-bp oligonucleotide containing an 18-bp random sequence^17^ was synthesized: 5′-GAGCCTCGTC TCCTGACTAT GCANNNGTNN NCTNNNAGNN NTGNNNCANN NTGCATCGTT GTGAGACGCA TGCTGCA-3′, along with the same reverse primer as above. Barcode extension, cloning, and plasmid preparation were performed as described. The full-length sequence is provided in Supplementary Table 3.

### Lentivirus production

HEK293T cells were seeded at 3 × 10^6^ cells per 10-cm dish. The following day, 2.5 μg of the lentiviral vector was mixed with 1.25 μg of pCMV-VSVG-RSV/Rev envelope vector and 1.25 μg of pMDLg/pRRE packaging vector in 250 μL Opti-MEM Reduced Serum Medium (Gibco, 31985-070). FuGENE HD Transfection Reagent (16 µL) (Promega, E2311) was added, the mixture was vortexed, left to incubate for 15 min, and carefully transferred to the HEK293T culture. The medium was refreshed 16–24 h post-transfection, and virus supernatant was collected 48 and 72 h later, passed through a 0.45 µm filter (Millipore Milex-HV, SLHVR33RS), aliquoted, snap-frozen in liquid nitrogen, and stored at −80°C until further use.

### Integrated multimodal experimental platform specialized in analyzing inter/intraclonal heterogeneity (IMPACH)

IMPACH, a versatile platform, combines single-cell-compatible barcoding with shRNA-mediated gene perturbation. By tuning the number of clones tracked (ranging from tens to hundreds), culturing strategies (pooled or isolated), and expansion time (i.e., number of cell divisions), IMPACH accommodates a broad range of applications, from monitoring clonal dynamics to quantifying inter- and intraclonal variability (**Extended Data Fig. 3**).

### IMPACH–Experimental procedure for interclonal analysis (Fig. 1)

Experimental parameters:

- Number of clones tracked: ∼400 clones per shRNA (∼2,000 clones in total)
- Culturing strategy: Pooled culture
- Expansion time: ∼15 cell divisions

To assess interclonal heterogeneity, BxPC3 cells were seeded at a density of 5,000 cells per well in 24- well plates. The following day, the cells were transduced with shPseuMO-Tag lentiviral vectors encoding shRNAs targeting KDM6A, SETD2, ARID1A, SMARCA4, or a nontargeting control (Scramble) in the presence of 10 µg/mL polybrene. The next day, 400 transduced BxPC3 cells, representing approximately 400 individual clones, were re-seeded into 24-well plates and cultured with 2 μg/mL puromycin for 3 days to select the successfully transduced cells. The cells were expanded until reaching approximately 10 million cells. After 19–26 days post-infection, the expanded populations were cryopreserved in Bambanker (Nippon Genetics) and stored until subsequent analyses. This procedure was performed independently for each shRNA and resulted in a total of approximately 2,000 clones analyzed across the five conditions.

### IMPACH–Experimental procedure for intraclonal analysis (Fig. 2, 3)

Experimental parameters:

- Number of clones tracked: 28
- Culturing strategy: Isolated single-cell cloning
- Expansion time: ∼20 cell divisions

To evaluate intraclonal heterogeneity, BxPC3 cells were seeded at a density of 5,000 cells per well in 24-well plates. The following day, the cells were transduced with shPseuMO-Tag lentiviral vectors encoding shRNAs targeting ARID1A (ARI3 and ARI4) or a nontargeting control (SCR2) in the presence of 10 µg/mL polybrene. The following day, the transduced cells were subjected to limiting dilution in 96- well plates to establish single-cell-derived clones and cultured with 2 μg/mL puromycin (3 days) to select cells infected with shPseuMO-Tag. Each clone was expanded until it reached approximately 1 million cells. The expanded clonal populations were cryopreserved in Bambanker (Nippon Genetics) and stored for subsequent assays. A total of 28 clones (8 SCR2 clones, 9 ARI3 clones, and 11 ARI4 clones) were used in subsequent experiments.

### *In vivo* cell-mixing metastasis assay

For the *in vivo* metastasis assay, 10 individual clonal populations (SCR2#01, SCR2#04, SCR2#05, SCR2#06, SCR2#13, ARI4#07, ARI4#14, ARI4#20, ARI4#21, and ARI4#25) were harvested and resuspended in Hank’s Balanced Salt Solution (Thermo Fisher Scientific). Equal numbers of cells from each clone were mixed, and a total of 20,000 cells in 200 μL were injected intravenously into the lateral tail vein of 7- to 8-week-old female NOD scid gamma (NSG) mice (Jackson Laboratory Japan). Lung tissues were collected at 14 or 28 days post-injection. Genomic DNA (gDNA) was extracted using the DNeasy Blood & Tissue Kit (Qiagen, 69504).

For re-barcoding experiments, two clones (SCR2#04 and ARI4#21) were independently re-barcoded, harvested, and resuspended in Hank’s Balanced Salt Solution. A total of 0.2 million cells in 200 μL were intravenously injected into the tail veins of 7-to 8-week-old female NSG mice. The remaining cells (∼1 million) were plated into 10-cm dishes in triplicate and cultured under standard conditions. Upon reaching confluence, the cells were passaged at a 1:5 ratio for five consecutive passages. Genomic DNA was extracted from the confluent cells after the fifth passage using the DNeasy Blood & Tissue Kit. All animal procedures were performed in accordance with protocols approved by the Animal Care and Use Committee of the Japanese Foundation for Cancer Research (approval number: 22-01-2).

For barcode library preparation, 500 ng of extracted gDNA was used as input for first-round PCR using the NEBNext High-Fidelity 2× PCR Master Mix (NEB, M0541). The thermal cycling conditions were as follows: 98°C for 30 s; followed by 19–25 cycles of 98°C for 10 s, 67°C for 30 s, and 72°C for 30 s. The PCR products were purified using 0.6× SPRIselect beads (Beckman Coulter) and eluted in 10 μL of TE buffer. The first-round PCR products were diluted 1:5 and used as templates for second-round PCR (50 μL reactions) to incorporate sample indices. The thermal cycling conditions were as follows: 98°C for 30 s, followed by 5 cycles of 98°C for 10 s and 72°C for 30 s. Dual-sided size selection was performed using SPRIselect beads with exclusion and selection ratios of 0.8× and 1.2×, respectively. The final products were eluted in 10 μL of TE buffer. The library size distribution and concentration were assessed using a TapeStation HSD1000 System (Agilent Technologies) and a Quantus Fluorometer (Promega), respectively. The libraries were sequenced on an Illumina NextSeq 550 platform using paired-end reads (Read 1: 75 bp; Index 1: 8 bp; Index 2: 8 bp; Read 2: 75 bp). The primer sequences used for library preparation are provided in Supplementary Table 4.

### scRNA-seq library preparation and sequencing

For the molecular profiling of clones analyzed using the IMPACH platform (**Figs. 1–3**), we performed single-plex, single-cell RNA sequencing to minimize batch effects. Barcoded cell populations transduced with shPseuMO-Tag vectors targeting KDM6A, SETD2, ARID1A, SMARCA4, or a nontargeting control (shSCR) were processed individually (**Fig. 1**), along with 28 distinct clones derived from shSCR2-, shARI3-, or shARI4-transduced cells (**Figs. 2 and 3**).

Cryopreserved cell stocks were rapidly thawed at 37°C, and cell viability was confirmed to exceed 90% before further processing. An equal number of viable cells was pooled and loaded onto a 10x Genomics Chromium controller for single-cell partitioning and barcoding. cDNA synthesis and library preparation were performed using the Chromium Single Cell 3′ v3 protocol (10x Genomics, Pleasanton, CA) based on the manufacturer’s instructions. Libraries were sequenced on an Illumina NextSeq 550 platform (Illumina, CA, USA) with paired-end reads (Read 1: 28 bp; Index 1: 10 bp; Index 2: 10 bp; Read 2: 90 bp). To analyze SMARCA4 inhibition by AU-15330 (**Fig. 4**), cells from distinct clones treated with DMSO or AU-15330 were pooled within each treatment group. Specifically, clones exposed to the same conditions were mixed in equal proportions before sample multiplexing using the 10x Genomics CellPlex 3′ Multiplexing Kit Set A. Equal numbers of pooled cells were loaded into the Chromium controller and processed as described above. Demultiplexing of individual clones was performed using a custom PseuMO-Tag Decoder pipeline (see below), whereas CellPlex sample identities were resolved using Cell Ranger v6.1.2.

To selectively enrich reads containing the PseuMO-Tag sequence, a three-round nested PCR strategy was used. For the first-round PCR, ∼20 ng of scRNA-seq library was amplified in a 100 μL reaction using NEBNext High-Fidelity 2× PCR Master Mix (NEB, M0541), and then split into two 50 μL reactions. The thermal cycling conditions were as follows: 98°C for 30 s; 15–20 cycles of 98°C for 10 s, 67°C for 30 s, and 72°C for 20 s. The pooled reactions were purified with SPRIselect beads (Beckman Coulter) using a dual size-selection (exclusion ratio: 0.56×; selection ratio: 0.85×) and eluted in 10 μL of TE buffer. For the second-round PCR, 10 ng of the purified product was used in a 50 μL reaction under the following conditions: 98°C for 30 s; 5 cycles of 98°C for 10 s, 65°C for 30 s, and 72°C for 20 s. The products were purified using 0.9× SPRIselect beads and eluted in 10 μL of TE buffer. For the third-round PCR, 10 ng of the second-round product was amplified using the same PCR conditions with 3 cycles at 67°C annealing. The final products were purified with 0.9× SPRIselect beads and eluted in low-EDTA TE buffer. The final barcode-enriched libraries were quality-checked using an Agilent TapeStation (HSD1000 kit) and quantified with a Quantus Fluorometer (Promega). Sequencing was done using an Illumina NextSeq 550 platform (paired-end, Read 1: 75 bp; Index 1: 8 bp; Index 2: 8 bp; Read 2: 75 bp). The primer sequences are listed in Supplementary Table 4.

### scATAC-seq library preparation and sequencing

For single-cell chromatin accessibility profiling, we performed single-plex, single-cell ATAC sequencing to minimize batch effects. Barcoded cell populations transduced with the shPseuMO-Tag vectors targeting KDM6A, SETD2, ARID1A, SMARCA4, or a nontargeting control (shSCR) were processed individually (**Fig. 1**). Similarly, the 28 clones derived from shSCR2-, shARI3-, or shARI4-transduced cells were processed (**Figs. 2 and 3**). The pooled clone mixtures were treated with either DMSO or AU-15330 (**Fig. 4**).

Cryopreserved cells were thawed at 37°C, and viability was confirmed to exceed 90%. Equivalent numbers of viable cells were used for each condition. Single-cell ATAC-seq libraries were generated using the SureCell ATAC-Seq Library Prep Kit and SureCell ddSEQ Index Kit (Bio-Rad) based on the manufacturer’s instructions. Libraries were sequenced on an Illumina NextSeq 550 platform using the following read structure: Read 1, 118 cycles (custom primer); i7 Index, 8 cycles; Read 2, 40 cycles. Raw FASTQ files were processed using the ATAC-Seq Analysis Toolkit (Bio-Rad) for debarcoding and alignment. Demultiplexing of individual clones was performed using a custom PseuMO-Tag Decoder pipeline.

To selectively enrich the reads containing the PseuMO-Tag barcode, two-round nested PCR was performed. For first-round PCR, ∼50 ng of scATAC-seq library DNA was used in a 100 μL reaction that was split into two 50 μL aliquots. The thermal cycling conditions were as follows: 98°C for 30 s; 15–20 cycles of 98°C for 10 s, 67°C for 15 s, and 72°C for 15 s. The pooled products were purified with SPRIselect beads (dual size-selection, exclusion: 0.95×, selection: 1.2×) and eluted in 10 μL TE buffer. The second-round PCR used 10 ng of the purified product in a 50 μL reaction, cycled at 98°C for 30 s, followed by 5 cycles of 98°C for 10 s, 66°C for 15 s, and 72°C for 15 s. The products were purified again using the same dual size-selection strategy. The final barcode-enriched libraries were assessed using an Agilent TapeStation (HSD1000 kit) and quantified using a Quantus Fluorometer (Promega). Sequencing was performed on an Illumina NextSeq 550 platform with the same read structure as the original scATAC-seq libraries. The primer sequences used for enrichment are listed in Supplementary Table 4.

### CUT&Tag library preparation and sequence

CUT&Tag was used to profile SMARCA4 binding in parental BxPC3 cells and in the clonal populations, SCR2#01, SCR2#04, ARI4#14, and ARI4#21. The experiments were performed using the CUT&Tag-IT Assay Kit (Active Motif, #53160) with modifications based on the manufacturer’s protocol and a previous report^55^. Approximately 200,000 cells were fixed in 0.1% paraformaldehyde for 10 min at room temperature and quenched with 125 mM glycine. After washing, the cells were gently scraped, incubated with concanavalin A-coated magnetic beads, and supplemented with 40 μL of CUT&Tag-IT Spike-in Nuclei (Active Motif, #53168) for normalization. The bead-bound nuclei were resuspended in 50 μL of antibody buffer containing 1 μL anti-SMARCA4 antibody (Brg1 [D1Q7F] Rabbit mAb, Cell Signaling Technology, #49360) and 1 μL of spike-in antibody (Active Motif, #53168). The samples were incubated overnight at 4°C on a nutator. After washing to remove unbound antibodies, guinea pig anti-rabbit secondary antibody (Active Motif, included in the kit) was added and incubated for 30 min at room temperature. After additional washes, the nuclei were incubated with CUT&Tag-IT-assembled pA-Tn5 Transposomes for 1 h at room temperature. Tagmentation was done by incubating the samples in Tagmentation Buffer for 2 min at 4°C, followed by 20 min at 37°C. To release tagmented DNA, the samples were treated with SDS and Proteinase K at 55°C for 1 h. DNA was purified using the DNA Purification Columns provided in the kit. Library amplification was achieved using indexed primers based on the manufacturer’s instructions. Libraries were size-selected using SPRIselect beads (Beckman Coulter), assessed for fragment size distribution with an Agilent TapeStation (HSD1000), and quantified using a Quantus Fluorometer (Promega). The final libraries were sequenced on an Illumina NextSeq 550 platform with paired-end reads (Read 1: 38 bp; Index 1: 8 bp; Index 2: 8 bp; Read 2: 38 bp).

### Transcriptomic heterogeneity estimation in the clinical samples

To determine the relationship between the expression of epigenetic modifiers^22^ and transcriptomic heterogeneity in primary human pancreatic tumors, we used two complementary computational approaches with bulk RNA-seq data from The Cancer Genome Atlas (TCGA) pancreatic adenocarcinoma (PAAD) cohort. First, we adopted Shannon’s equitability-based method, with minor modifications from a previously published protocol by Hinohara *et al.*^23^. For each epigenetic modifier, the patients were stratified into four quartile groups of equal size based on gene expression levels. Shannon’s equitability was calculated for each tumor based on the distribution of expression values across all detected genes. This metric normalizes Shannon’s entropy to a [0,1] scale, in which 1 indicates uniform gene expression and values near 0 indicate transcriptional dominance by a small subset of genes.

Second, we used a PCA-based transcriptomic diversity scoring method adapted from Ma *et al.*^24^. Principal component analysis (PCA) was performed using the prcomp function in R for all patient samples. Top *n* principal components (PCs) were extracted to represent the dominant axes of transcriptomic variation. Each patient was projected into this PC space, and the centroid of the population was computed for each quartile group. Transcriptomic diversity was defined as the mean Euclidean distance of each patient to the global centroid and calculated as follows:

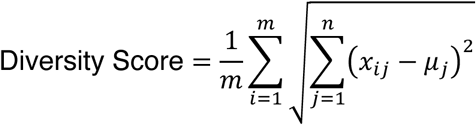

Here, *m* is the number of patients, *n* is the number of PCs, *x_ij_* is the *j*th PC score of the *i*th patient, and *μ_j_* is the mean of the *j*th PC for all patients. To minimize the influence of outliers, samples with PC values exceeding ±3 standard deviations for any of the top three PCs were excluded. The top 30 PCs were selected based on the cumulative variance explained and the stability of the resulting diversity scores.

### Survival analysis

Survival analysis was performed using the survival and survminer packages in R, based on TCGA PAAD patients with available clinical follow-up data. Kaplan–Meier survival curves were generated by stratifying patients according to the average expression of GEP3-associated genes. Statistical significance between the groups was assessed using the log-rank test.

### Analysis of the clonal distribution

To extract high-quality barcode sequences from the raw amplicon sequencing reads, the 5′ adaptor sequences (5′-GTCGAGGCAG GAAACAGCTA TGACTATGCA-3′) and 3′ adaptor sequences (5′-TGCATCGTTG AGCAATAA-3′) were trimmed using cutadapt (v4.4). Reads containing ambiguous nucleotides (Ns) shorter than 28 nucleotides or lacking the characteristic barcode motif (5′-NNNGTNNNCT NNNAGNNNTG NNNCANNN-3′) were excluded. The remaining reads were filtered to retain only those with an average Phred quality score ≥30. The qualified barcode sequences were subjected to error correction using Starcode (v1.4) with a maximum Levenshtein distance of 1. The corrected barcode counts were aggregated for each sample and subsequently used for downstream clonal distribution analyses.

### IMPACH–Clone calling for inter- and intraclonal heterogeneity; PseuMO-Tag Decoder

Demultiplexing of individual clones was done using a custom pipeline, *PseuMO-Tag Decoder*, which was developed to assign clonal identities based on synthetic barcodes introduced via the shPseuMO-Tag vector (**Extended Data Fig. S2**).

For scRNA-seq data, amplicon-seq FASTQ files were generated as described in the “scRNA-seq library preparation and sequencing” section. From Read 1, the first 16 bp and 12 bp were extracted and designated as the cell barcode (CB) and unique molecular identifier (UMI), respectively. Error correction was performed and allowed up to one mismatch (Hamming distance ≤ 1). Adapter sequences (5′-CAGGAAACAGCTATGACT-3′) were trimmed from Read 2 using cutadapt, and a barcode region composed of HashTag–PseuTag–HashTag (5 bp–10 bp–5 bp) was extracted. The HashTag and PseuTag sequences were independently error-corrected, allowing one mismatch. PseuTag error correction was performed using Starcode^56^. To remove PCR duplicates, the read counts were collapsed by UMI to generate unique CB–PseuTag pairs. Only cells with at least two UMI-supported PseuTag reads were retained for downstream analysis.

For scATAC-seq data, clonal assignment was done using the corresponding amplicon-seq FASTQ files (see “scATAC-seq library preparation and sequencing” section). CBs were identified from Read 1 using the ATAC-Seq Analysis Toolkit (Bio-Rad). Read 2 was processed similarly to the scRNA-seq workflow using the adapter sequence: 5′-CGCTCGCTAGTTATTGCTCA ACG-3′, followed by HashTag and PseuTag extraction and error correction as described above.

To integrate clonal information across the scRNA-seq and scATAC-seq datasets, shRNA identities were assigned to cells, in which a single HashTag accounted for >60% of the total UMI counts. For clonal classification, a cell-by-PseuTag count matrix was constructed separately for each shRNA group.

Hierarchical clustering was performed using the heatmap3 R package (v1.1.9) with Ward.D linkage and cells within the same cluster were assigned to the same clone. In Figure 4, cells were directly assigned to individual clones if a single dominant PseuTag accounted for more than 60% of the total PseuTag counts. This framework enabled the robust linkage of single-CBs to their corresponding clonal identities (PseuTags), which facilitated clone-aware single-cell transcriptomic and epigenomic analyses.

### IMPACH–Data analysis for interclonal heterogeneity (Fig. 1)

For scRNA-seq data, raw sequencing reads were processed using Cell Ranger (v6.1.2, 10x Genomics) with default parameters and aligned to the GRCh38 human reference genome to generate gene-by-cell expression matrices. Demultiplexing of individual clones was performed using the PseuMO-Tag Decoder pipeline (see above). Downstream analysis was done using the Seurat R package (v4.0.1)^57^. Low-quality cells were filtered out based on the following criteria: mitochondrial transcript proportion of >20%, <200 detected genes, or >4,000 detected genes. Gene expression matrices were normalized using Seurat’s NormalizeData() function, which performs log-normalization by scaling each UMI count for each cell to 10,000, followed by natural logarithmic transformation. Data were subsequently scaled using ScaleData(). Highly variable genes were identified using FindVariableFeatures() with the “vst” method, and the top 2,000 genes were selected. PCA was performed with RunPCA(). For dimensionality reduction and clustering, the top 30 PCs were used as input for FindNeighbors() and FindClusters() with a resolution of 0.4. UMAP visualization was generated using RunUMAP() based on the same PCs. Knockdown efficiency of the target genes was evaluated by calculating the average expression values for each shRNA group using AverageExpression(), and visualized with the ComplexHeatmap package (v2.6.2). Moreover, differentially expressed genes (DEGs) for each clone were identified using Seurat’s FindMarkers() function. To ensure robust detection, an equal number of cells were randomly subsampled 100 times from the scramble control group to match the number of cells per clone. For each iteration, log₂ fold changes and adjusted P-values were calculated. Genes with a median log₂ fold-change of ≥0.25 and a median adjusted P-value of ≤0.1 across iterations were defined as DEGs for that clone. Functional enrichment analysis of the DEGs was done using the Enrichr web server^58–60^, and the enriched pathways were visualized using Appyter available through Enrichr.

Regulatory network inference of MYC-associated transcription factors (TFs) was performed using the decoupleR package (v1.6.1)^61^, based on the CollecTRI prior knowledge network. TF activity scores were inferred per cell using the univariate linear model (ULM) method implemented in run_ulm(), and extracted with get_acts() for downstream analyses. Regulatory network structures were visualized using plot_network(), specifying selected MYC-related TFs as sources and limiting visualization to the top 45 target genes. The positive and negative regulatory edges were colored dark green and dark red, respectively.

For scATAC-seq data, raw reads were processed using the ArchR package (v1.0.2)^62^ with modifications based on previously described workflows^49, 63^. Demultiplexing of individual clones was done using the PseuMO-Tag Decoder pipeline (see above). Analyses were conducted using the hg19 human genome assembly specified with addArchRGenome (“hg19”). Initial quality control was performed with createArrowFiles() to compute the transcription start site (TSS) enrichment scores and fragment counts per nucleus. Cells with TSS enrichment scores <8 or fewer than 2,500 unique fragments were excluded. Potential doublets were removed using addDoubletScores() with parameters k=10, knnMethod=“UMAP”, and LSIMethod=1. Dimensionality reduction was done using addIterativeLSI() based on genome-wide 500-bp tiles. Clustering was performed using addClusters() and Seurat’s FindClusters() function, and the UMAP projection was generated by addUMAP(). To identify reproducible peak regions, pseudo-bulk replicates were generated using addGroupCoverages(), and peak calling was conducted with MACS2 (v2.2.7.1) through addReproduciblePeakSet() with parameters shift=−40, extsize=80, --nomodel, and --nolambda. Resulting peak set was incorporated into the project using addPeakMatrix(). Marker peaks were identified using getMarkerFeatures() and filtered with getMarkers() under the thresholds Pval of ≤0.05 and Log2FC of ≥1. TF activity was inferred using chromVAR deviation scores. Motif annotations were added with addMotifAnnotations() using the CIS-BP database, and putative regulatory TFs were inferred based on the overlap between the motif locations and the marker peak regions.

### IMPACH–Data analysis for intraclonal heterogeneity (Figs. 2 and 3)

For scRNA-seq analysis, raw sequencing reads were processed using the Cell Ranger pipeline (v6.1.2, 10x Genomics) and aligned to the GRCh38 human reference genome to generate gene-by-cell expression matrices as described above (see IMPACH–Data analysis for interclonal heterogeneity). Demultiplexing of individual clones was done using the custom PseuMO-Tag Decoder pipeline.

Downstream analyses were performed using the Seurat R package (v4.0.1)^57^ with modified quality control thresholds. Cells with >10% mitochondrial transcript content, fewer than 800 detected genes, or more than 4,000 detected genes were excluded.

Gene expression matrices were log-normalized and scaled, and PCA was performed as previously described. The top 30 PCs were used for clustering (FindNeighbors() and FindClusters() with a resolution of 0.4) and UMAP visualization.

Gene expression programs (GEPs) were identified using consensus non-negative matrix factorization (cNMF)^41^. Factorization was performed with seven components (--components 7) and a local density threshold of 0.2. The top 50 genes for each GEP were subjected to enrichment analysis using the Enrichr web server^58–60^ against the MSigDB Hallmark 2020 and GO Biological Process 2025 databases. GEPs were manually annotated based on enrichment profiles. Clone-level GEP annotations were assigned by computing the median GEP activity per clone, and hierarchical clustering was performed using the heatmap3 package (method = “ward.D2”). Clones were grouped into four transcriptional classes based on the dendrogram structure. Ternary plots of GEP1, GEP3, and GEP4 activity scores were generated for the ARID1A knockdown clones using the ggtern package (v3.5.0).

The transcriptional regulatory inference of GEP3-associated programs was performed with decoupleR (v1.6.1)^61^, and leveraged the CollecTRI database for prior knowledge. TF activity scores were inferred by the ULM method (run_ulm()).

For the scATAC-seq data, the raw reads were processed and analyzed using the ArchR package (v1.0.2)^62^ as described above, with minor modifications. Demultiplexing was performed using the PseuMO-Tag Decoder pipeline. Clone-specific marker peaks were identified with getMarkerFeatures() followed by filtering with thresholds of FDR ≤0.5 and log₂ fold-change ≥0.25. TF activity was inferred using chromVAR deviation scores with motif annotations from the CIS-BP database. Top 100 most variable motifs (z-scored) were visualized using the ComplexHeatmap package (v2.6.2).

To quantify chromatin accessibility at GEP3-associated regulatory regions, 100 cells per clone were randomly sampled, and the number of Tn5 fragments overlapping the promoter regions of the top five GEP3-associated genes was calculated using getFragmentsFromProject(). Sampling was repeated 100 times per clone, and the results were visualized using boxplots (median, interquartile range (IQR), and whiskers at 1.5× IQR).

### IMPACH–Data analysis for drug treatment (Fig. 4)

For scRNA-seq analysis, data processing, clone demultiplexing (PseuMO-Tag Decoder), and CellPlex sample assignment (Cell Ranger v6.1.2) were performed as described above. Cells with >20% mitochondrial transcript content, fewer than 200 detected genes, or more than 4,000 detected genes were excluded. Normalization, scaling, PCA, clustering, and UMAP visualization followed the procedures described earlier. The GEP3 module score was calculated for each cell using the AddModuleScore() function in Seurat.

For the scATAC-seq data, processing and analysis were conducted using the ArchR package (v1.0.2)^62^, following the same workflow described above. Demultiplexing was conducted using the PseuMO-Tag Decoder pipeline. Quality control, dimensionality reduction (LSI), clustering, UMAP projection, peak calling using MACS2, and the addition of peak matrices were performed as previously described.

### CUT&Tag data processing

Adapter-trimmed, paired-end reads were aligned to the human reference genome (hg38) using Bowtie2 (v2.4.5) with the parameters: --very-sensitive -X 2000 --no-mixed --no-discordant. The resulting SAM files were converted to BAM format and sorted using SAMtools (v1.15.1). To retain only uniquely mapped reads, the alignments containing the “XS” tag were excluded. PCR duplicates were removed using the MarkDuplicates tool from Picard (v2.27.5) and the options VALIDATION_STRINGENCY=LENIENT and REMOVE_DUPLICATES=true. Reads mapping to genomic blacklist regions (ENCODE hg38 blacklist v2) were removed using intersectBed from the Bedtools suite (v2.30.0) with the corresponding blacklist BED file. Peak calling was performed using MACS2 (v2.2.7.1) in narrowPeak mode with the parameters: --format BAMPE and --qvalue 0.1. To normalize for spike-in controls, reads were also aligned to the *Drosophila melanogaster* reference genome (Dmel_A4_1.0) using the same alignment pipeline. Sample-specific scaling factors were calculated based on mapping percentages. Quantification of the read abundance around the peak centers (±2 kb) identified from the scATAC-seq data was performed using the computeMatrix function of the deepTools package (v3.5.1)^64^, and applying spike-in–normalized BAM files. The profiles were stratified by genomic region categories (promoter, intronic, exonic, and distal) and visualized using the plotProfile function in deepTools. Peak correlations between samples were calculated using the DiffBind package (v3.12.10), and peak annotations were obtained using the ChIPseeker package (v1.38.0).

### Quantification of the transcriptomic heterogeneity

To determine the effects of epigenetic modifier loss on transcriptomic heterogeneity, we calculated the pairwise normalized mutual information (NMI) scores (**Fig. 1d**; **Fig. 2d,e**) based on a method previously described by Marjanovic *et al*.^65^ and Fennell *et al*^5^.

Gene expression values for each clone or condition were discretized into 10 bins using either differentially expressed genes or NMF-derived signature genes as the input feature set. The entropy of an individual cell *X* was defined as follows:

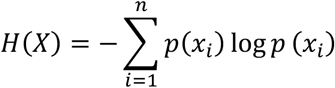

Here, *p*(*x_i_*) represents the empirical probability of the expression level of a gene falling into the *i*th bin. To capture the information shared between cell pairs, mutual information was calculated as follows:

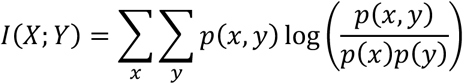

Here, *p*(*x*,*y*) denotes the joint distribution of binned gene expression across two cells, and *p*(*x*) and *p*(*y*) are their respective marginal distributions. The NMI between the two cells was then computed as follows:

To robustly estimate the population-level heterogeneity, 100 cells were randomly subsampled without replacement from each group, and the average pairwise NMI was computed across all cell pairs. This process was repeated 1,000 times to minimize sampling bias. NMI distributions were visualized using boxplots, in which the boxes represent the IQR, the center lines indicate the median, and the whiskers extend to 1.5 times the IQR.

To assess interclonal transcriptomic heterogeneity from scRNA-seq data (**Fig. 1g**), we computed a diversity score following the method described by Ma *et al*.^24^, with minor modifications. Briefly, PCA was performed on all high-quality cells to reduce dimensionality and extract the major sources of variation. PCA was conducted using the RunPCA function in the Seurat package (v4.0.1) with default parameters. Furthermore, each cell was represented as a vector in the space of the top *n* PCs, and the centroid of the population was defined as the mean vector across all cells. Diversity score was calculated as the average Euclidean distance from each cell to the centroid using the following equation:

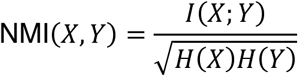

Here, *m* is the number of cells, *n* is the number of PCs, *x_ij_* is the value of the *j*th PC in the *i*th cell, and *μ_j_* is the mean of the *j*th PC across all cells. To minimize the effect of extreme outliers, cells with values exceeding ±3 standard deviations from the population mean in any of the first three PCs were excluded. Diversity scores were calculated separately for each clone and visualized as boxplots. In the boxplots, the lower and upper hinges represent the first and third quartiles, respectively, whereas the horizontal line within the box represents the median. The upper and lower whiskers extend to the highest and lowest values within 1.5 times the interquartile range (IQR) from the hinge.

For the interclonal analysis, the top 30 PCs were used for the diversity score calculation. Results were robust to the number of PCs used, with minimal variation observed when adjusting the number of components.

### Quantification of chromatin accessibility heterogeneity in the scATAC-seq data

To quantify the heterogeneity in chromatin accessibility, we calculated the peak entropy for each candidate cis-regulatory element (cCRE). A reproducible peak set was first obtained from the scATAC-seq data using the getMatrixFromProject() function in the ArchR package. Regarding functionally relevant regulatory elements, peaks were filtered to retain only those overlapping with ENCODE candidate cis-regulatory elements (cCREs). ENCODE cCRE annotations were downloaded from the UCSC Genome Browser (GRCh38/hg38) and converted to GRCh37/hg19 coordinates using the UCSC LiftOver tool. Tn5 insertion sites were inferred from aligned reads by correcting strand-specific bias: +4 bp for reads mapped to the plus strand and −5 bp for reads on the minus strand^66, 67^. Moreover, downstream analyses were restricted to cCRE-overlapping peaks detected in at least 1,000 single cells. To calculate peak entropy, we adapted the single-cell entropy calculation described by Pastore *et al*.^33^. For each peak, the fraction of positive cells (fpc), defined as the proportion of single cells with at least one Tn5 insertion at that peak, was used to calculate the Shannon entropy as follows:

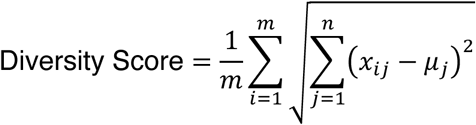

This metric captures the degree of dispersion in chromatin accessibility across the cell population at each cCRE. Peaks with intermediate fpc values (closer to 0.5) yield higher entropy, indicating greater heterogeneity, whereas those with values near 0 or 1 yield lower entropy, reflecting more uniform accessibility.

### Statistical analysis

Statistical analyses were conducted using R software. Parametric tests included unpaired two-tailed Student’s *t*-tests and one-way analysis of variance (ANOVA), followed by Dunnett’s or Tukey’s post hoc multiple comparison tests where appropriate. For nonparametric comparisons, the Wilcoxon rank-sum test was used. A *P*-value <0.05 was considered statistically significant.

### Code availability

Processed single-cell objects and custom PseuMO-Tag Decoder to reproduce analyses and figures is available at https://github.com/KMiyata7/PseuMO-Tag.

## Acknowledgements

We thank Dr. Kornelia Polyak (Dana-Farber Cancer Institute) for her insightful advice. We are also grateful to Drs. Shinji Nakaoka and Ryo Yamaguchi (Hokkaido University), and Drs. Naotoshi Nakamura and Shingo Iwami (Nagoya University) for their valuable suggestions regarding the mathematical analyses. We thank all members of the Maruyama laboratory for helpful discussions and support. This work was supported in part by JSPS KAKENHI Grant Numbers 23K18247 (to K.M.), 23K27446 (to K.M.), 24K02312 (to R.M.), the Japan Agency for Medical Research and Development (AMED) JP24ama221606 (to R.M.), the Kato Memorial Bioscience Foundation, the Uehara Memorial Foundation, the Takeda Science Foundation, and the Mochida Memorial Foundation for Medical and Pharmaceutical Research.

## Author contributions

K.M. and R.M. conceived the study and wrote the manuscript. K.M. performed most of the experiments with assistance of L.Y., Y.Y., K.K.. All authors approved the manuscript.

## Supplementary Information

Supplementary Table 1. Comparison of IMPACH using the shPseuMO-Tag with previously reported cell barcoding and perturbation systems. Summary table of the key features across different systems, including compatibility with genetic perturbations, clonal tracking resolution, and integration with single-cell ATAC-seq and RNA-seq. Integrated multimodal experimental platform specialized in analyzing inter/intraclonal heterogeneity (IMPACH) provides broader functional coverage and multimodal compatibility compared with previously established platforms.

Supplementary Table 2. List of genes that contain seven GEPs.

Supplementary Table 3. Sequence of shPseuMO-Tag and PseuMO-Tag. Supplementary Table 4. List of primers used in the manuscript.

## Extended Data

**Extended Data Fig. 1.**
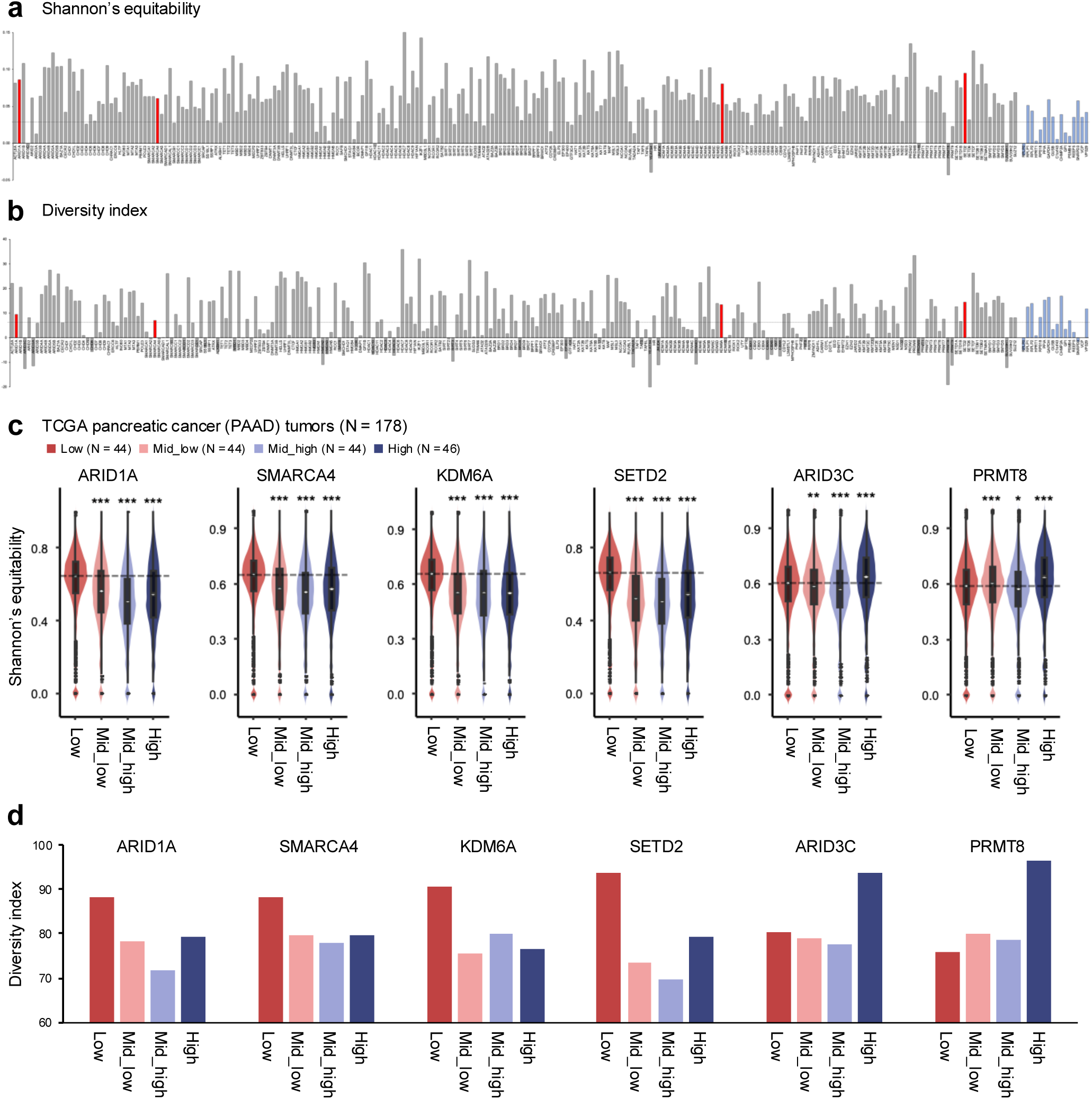
Association between epigenetic modifier expression and transcriptomic heterogeneity in pancreatic cancer clinical samples. (**a, b**) Transcriptomic heterogeneity was quantified in TCGA pancreatic adenocarcinoma (PAAD) tumors (N = 178) using either Shannon’s equitability index (**a**) or a principal component-based diversity index (**b**). Tumors were stratified into four expression quartiles: Low (N = 44), Mid-low (N = 44), Mid-high (N = 44), and High (N = 46), based on the expression of ARID1A, SMARCA4, KDM6A, SETD2, ARID3C, and PRMT8. The differences in heterogeneity between tumors with the highest and lowest expression levels for each gene are shown; dashed line indicates the median of housekeeping genes. Red bars represent selected epigenetic modifiers, while blue bars indicate housekeeping genes. (**c, d**) Violin plots of Shannon’s equitability index (**c**) and bar plots of the principal component-based diversity index (**d**) for tumors stratified by expression of the six indicated genes. The quartile groups are color-coded as Low, Mid-low, Mid-high, and High. *P* < 0.05, \*\**P* < 0.01, \*\*\**P* < 0.001 by one-way ANOVA followed by Dunnett’s post hoc multiple comparisons test.

**Extended Data Fig. 2.**
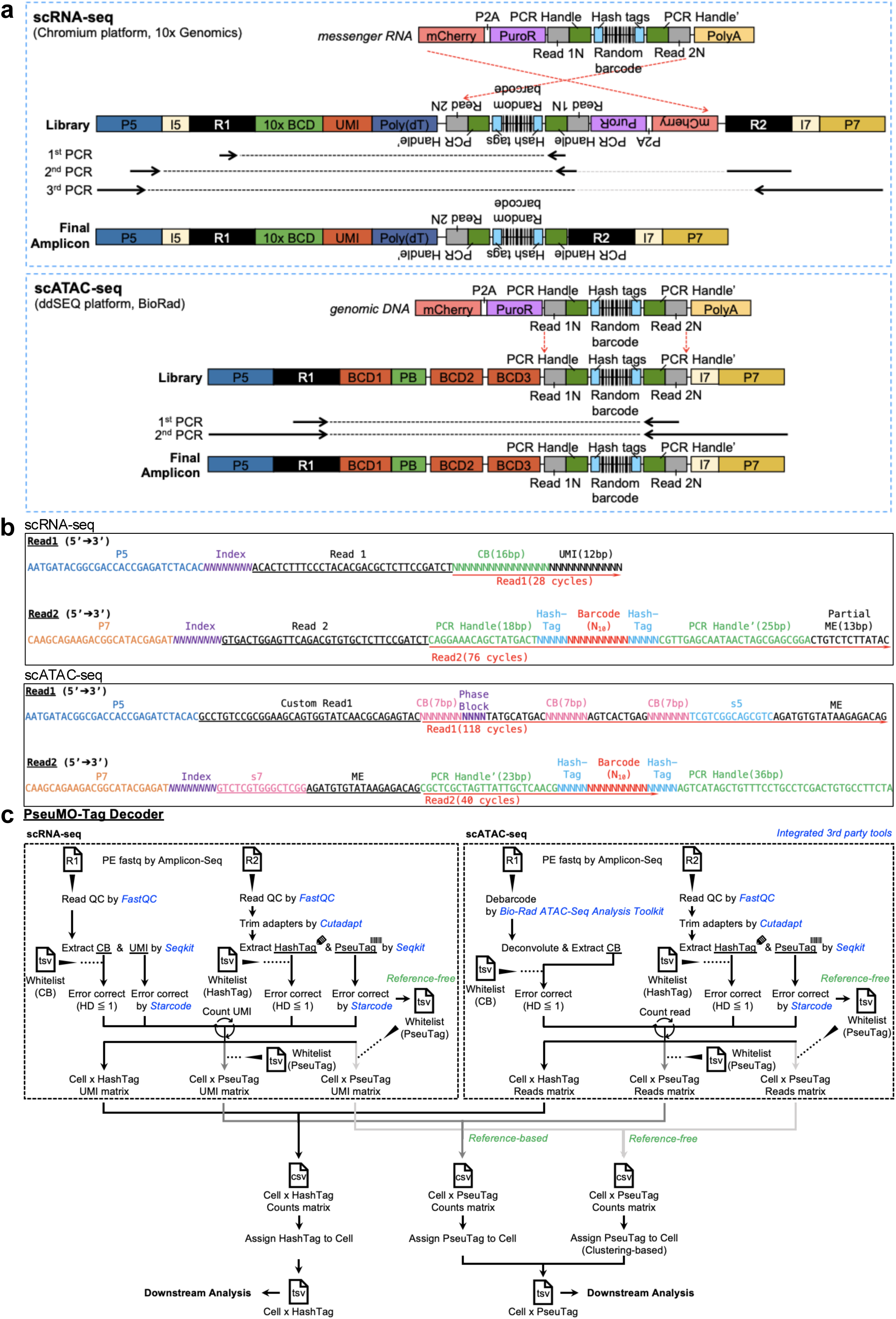
Overview of the PseuMO-Tag sequencing strategy and decoding pipeline. (**a**) Schematic of the library preparation workflows for PseuMO-Tag–integrated scRNA-seq (top) and scATAC-seq (bottom). scRNA-seq workflow follows the Chromium platform (10x Genomics) and incorporates shPseuMO-Tag transcripts into messenger RNA libraries. The scATAC-seq workflow is based on the ddSEQ platform (Bio-Rad) and enables the integration of shPseuMO-Tag barcodes into chromatin accessibility libraries for clonal tracking. (**b**) Read structures of scRNA-seq and scATAC-seq libraries generated using the PseuMO-Tag system, highlighting barcode positioning and sequencing configurations. (**c**) Schematic of the *PseuMO-Tag Decoder* pipeline for scRNA-seq and scATAC-seq data. Amplicon-seq reads generated from the respective libraries were processed to associate single-cell barcodes with clonal barcodes (PseuMO-Tags). The resulting matrices link each cell to its clonal identity through either reference-based or reference-free assignment, enabling downstream analyses.

**Extended Data Fig. 3.**
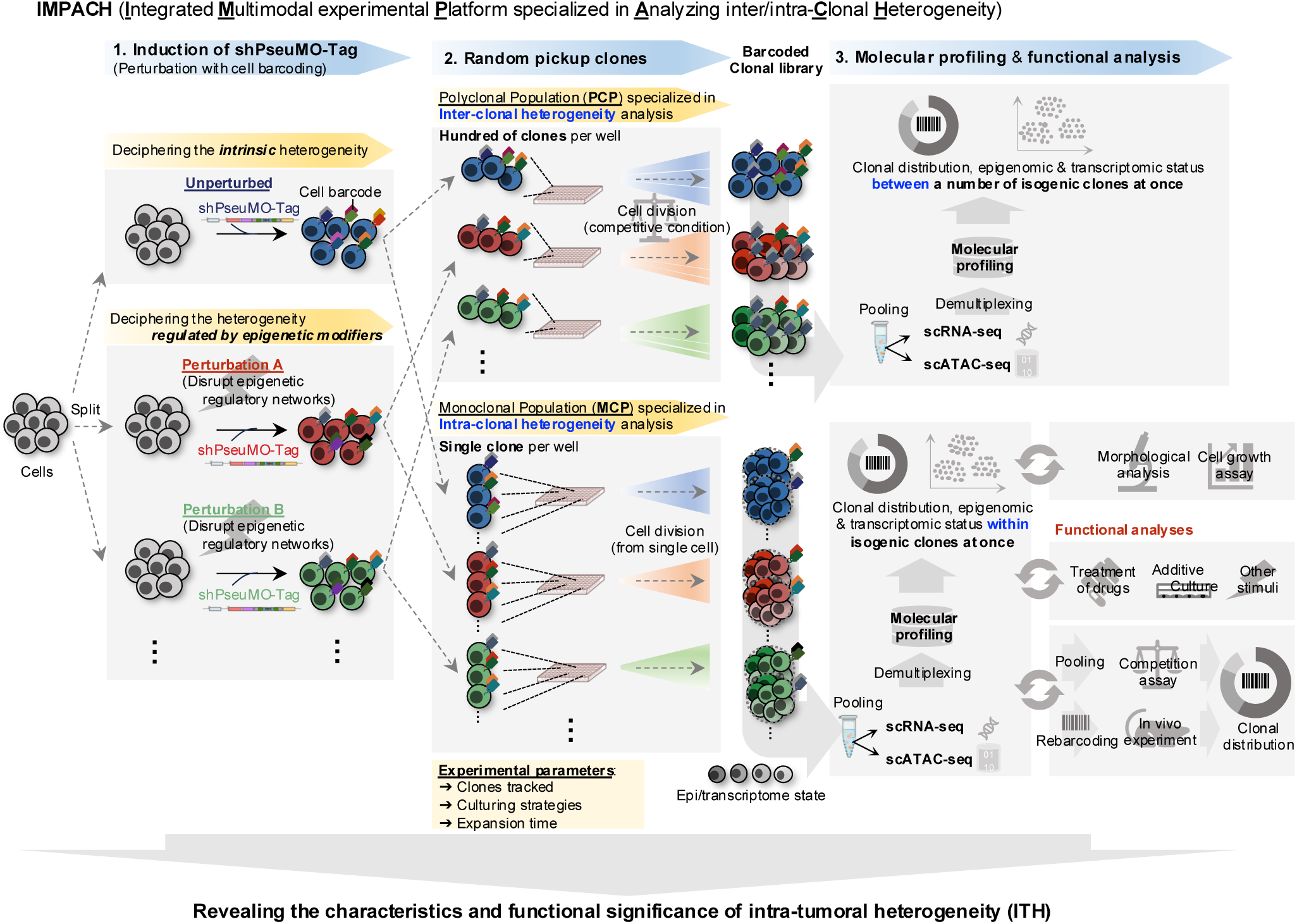
Schematic overview of the IMPACH system for decoding inter- and intraclonal heterogeneity. Schematic of the workflow of Integrated multimodal experimental platform specialized in analyzing inter/intraclonal heterogeneity (IMPACH). The platform integrates perturbation, clonal tracking, single-cell profiling, and functional interrogation to systematically investigate clonal heterogeneity. Step 1: Induction of the shPseuMO-Tag for barcoding and perturbation. The cells were split into multiple experimental arms and transduced with shPseuMO-Tag vectors. Unperturbed conditions enable the study of intrinsic heterogeneity, while perturbation conditions enable the assessment of how the disruption of targeted regulatory networks affects heterogeneity. Step 2: Clone isolation and expansion. Transduced cells were expanded using two culture strategies: (i) polyclonal populations (PCPs) containing hundreds of clones per well for analyzing interclonal heterogeneity; and (ii) monoclonal populations (MCPs) with one clone per well for analyzing intraclonal heterogeneity. By tuning experimental parameters, including the number of clones tracked (from tens to hundreds), culturing strategies (pooled versus isolated), and expansion time (i.e., cell division cycles), IMPACH supports a wide range of applications, from monitoring clonal dynamics to quantifying intraclonal variability. Step 3: Molecular profiling and functional analyses. Barcoded clonal libraries were pooled and demultiplexed using the PseuMO-Tag Decoder pipeline, enabling integrated scRNA-seq and scATAC-seq analysis. Molecular profiling assesses clonal diversity and epigenomic/transcriptomic states. Functional assays, including drug treatments, cell growth assays, re-barcoding experiments, and *in vivo* competition assays, are conducted to determine the phenotypic significance of clone-specific states. IMPACH system enables the parallel, high-throughput characterization of both inter- and intraclonal heterogeneity and their functional consequences, facilitating the dissection of the molecular logic underlying intratumoral heterogeneity (ITH).

**Extended Data Fig. 4.**
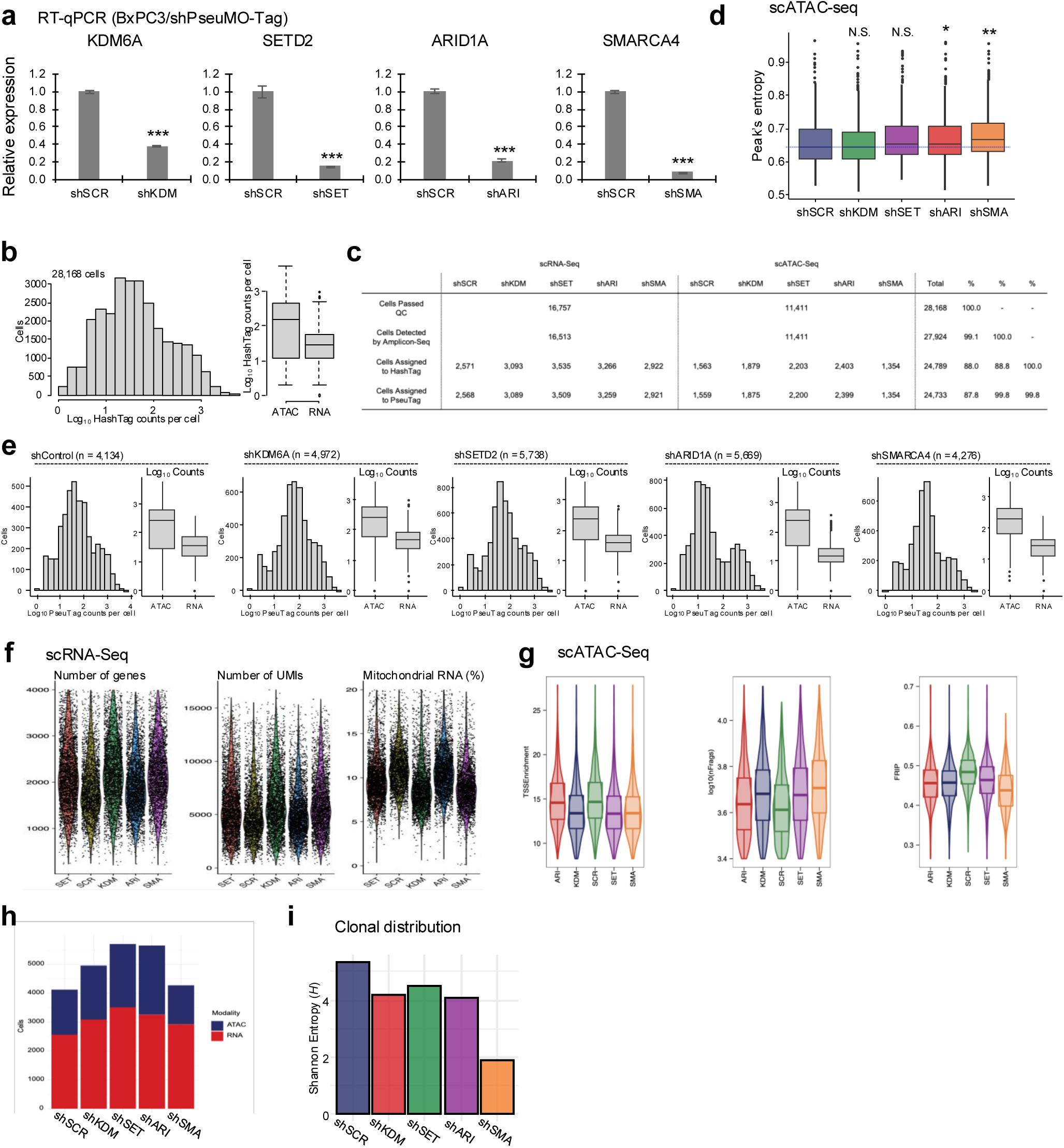
Demultiplexing performance and quality assessment of the IMPACH platform. (**a**) RT-qPCR analysis of KDM6A, SETD2, ARID1A, and SMARCA4 expression in BxPC3 cells transduced with shPseuMO-Tag constructs targeting each gene or a nontargeting control (SCR). Data represent the mean ± s.d. from biological triplicates. \*\*\**P* < 0.001 by two-tailed Student’s *t*-test. (**b**) Distribution of PseuMO-Tag counts per cell detected in scRNA-seq and scATAC-seq datasets (n = 28,168 total cells). Histogram (left) and box plot of log₁₀-transformed counts (right) illustrate barcode complexity across modalities. (**c**) Summary table of single-cell sequencing metrics. The number and proportion of cells passing quality control, as detected by Amplicon-seq, and successfully assigned to hash tags (shRNAs) and PseuMO-Tags are shown for each shRNA condition and modality. (**d**) Box plots of peak entropy for the shRNA conditions, calculated from chromatin accessibility at candidate cis-regulatory elements (cCREs). \**P* <0.05, \*\**P* <0.001, N.S., not significant by one-way ANOVA followed by Dunnett’s post hoc test. (**e**) Distribution of log₁₀-transformed PseuMO-Tag counts per cell across five shRNA conditions (shScramble, shKDM6A, shSETD2, shARID1A, shSMARCA4) in the scRNA-seq and scATAC-seq datasets. (**f**) Violin plots of single-cell RNA-seq quality metrics per shRNA condition, including the number of RNA molecules (nCount), number of detected genes (nFeature), and mitochondrial transcript fraction (%Mito). (**g**) Violin plots of single-cell ATAC-seq quality metrics per shRNA condition, including the number of fragments (nFrags), transcription start site (TSS) enrichment, and fraction of reads in peaks (FRiP). (**h**) Stacked bar plots showing the modality of cells detected (ATAC or RNA) across shRNA conditions. (**i**) Bar plot of Shannon entropy representing clonal distribution across shRNA conditions.

**Extended Data Fig. 5.**
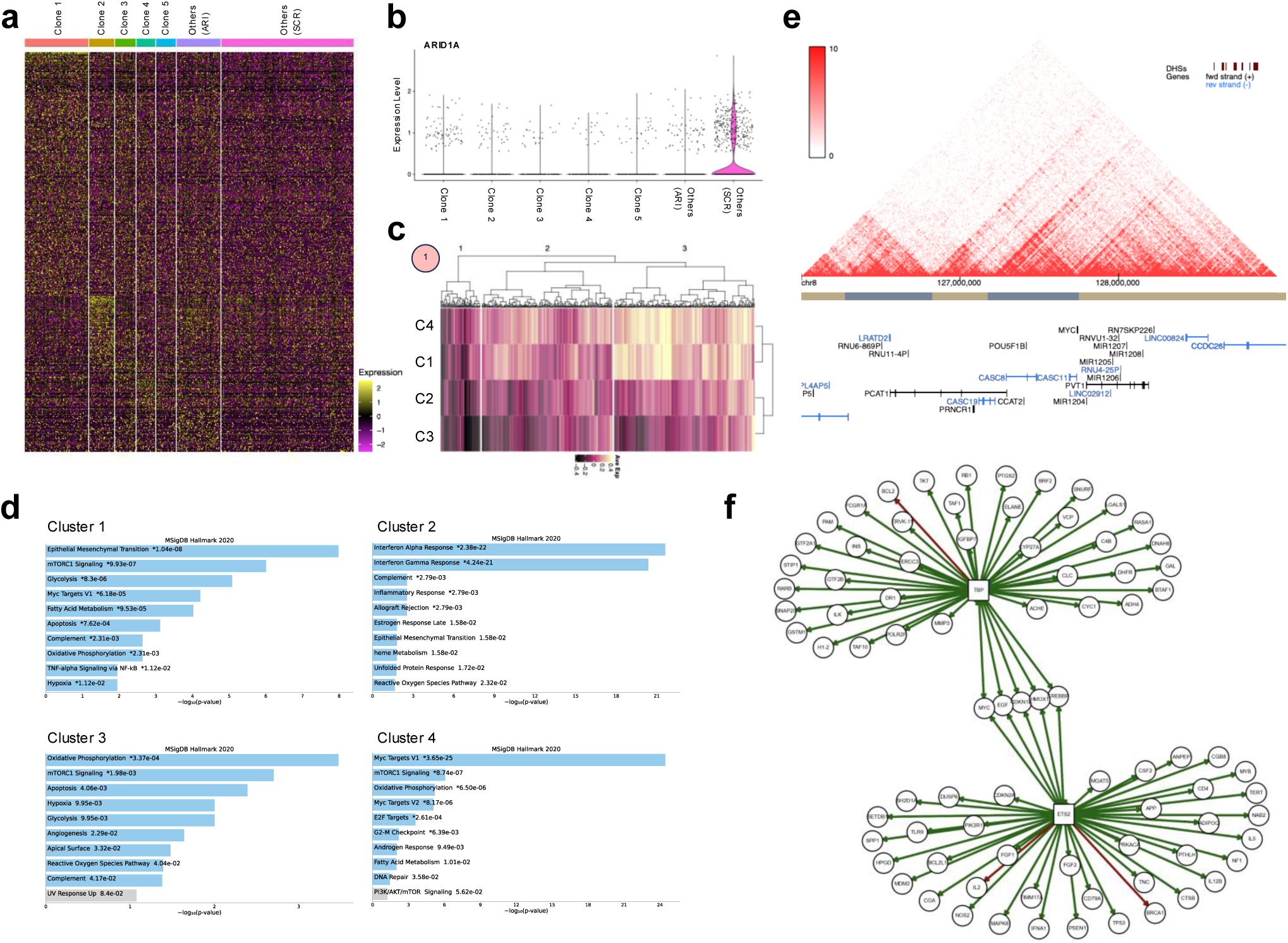
Pseudomultiomic analysis of ARID1A-deficient clones reveals MYC- and EMT-associated transcriptional reprogramming. (**a**) Heatmap showing the scaled expression of 525 differentially expressed genes across the top five ARID1A-knockdown clones, other ARID1A-knockdown clones, and scramble controls (SCR). (**b**) Violin plot displaying scaled ARID1A expression levels across individual clones corresponding to those shown in (a). (**c**) Heatmap showing the average expression of genes in each transcriptomic cluster (C1–C4), as defined by k-means clustering shown in Fig. 1j, within the dominant ARID1A-knockdown clone. (**d**) Gene set enrichment analysis for each transcriptomic cluster (C1–C4) based on hallmark gene sets from MSigDB (2020), performed using Enrichr. The top pathways are shown, ranked by adjusted *P*-value. (**e**) Chromatin conformation landscape around the MYC locus in BxPC3 cells, visualized using Hi-C data from the 3D Genome Browser^68, 69^ (hg38; chr8:126,000,000–129,000,000). (**f**) Regulatory network of MYC-related transcription factors identified in the dominant clone. The green arrows represent positive regulation, and red arrows represent negative regulation.

**Extended Data Fig. 6.**
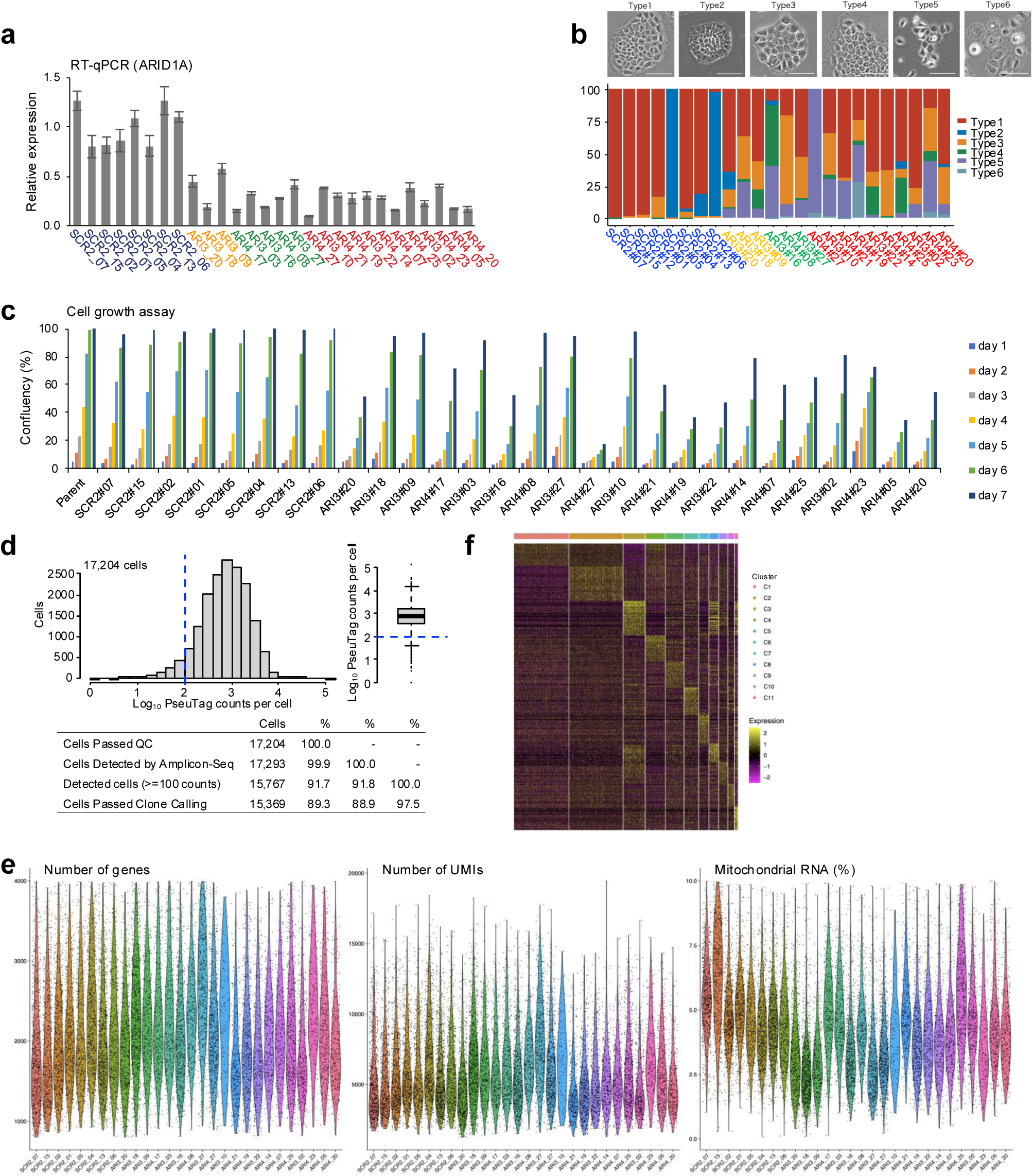
Characterization and quality metrics of single-cell RNA-seq data across 28 clones analyzed by IMPACH. (**a**) RT-qPCR analysis of ARID1A expression across 28 clones analyzed by IMPACH. Data represent the mean ± s.d. from biological triplicates. Expression levels were normalized to ACTB and shown relative to the average of the control (SCR2) clones. (**b**) Representative phase-contrast images of clonal morphologies classified into six types (Type 1–6). The bar plot shows the relative proportion of each morphology type across the clones. Scale bar, 35 µm. (**c**) Cell growth assay showing confluency percentages over 7 days for each clone. Measurements were taken daily using the IncuCyte live-cell imaging system. (**d**) Distribution of log₁₀-transformed PseuMO-Tag counts per cell across 17,204 single cells passing quality control. The box plot (right) and summary table (bottom) show the proportion of cells detected by Amplicon-seq, assigned to clones, and retained after clone calling. (**e**) Violin plots of single-cell RNA-seq quality metrics, including the number of detected genes (left), number of unique molecular identifiers (UMIs; center), and percentage of mitochondrial RNA (right), stratified by clone identity. (**f**) Heatmap of scaled expression levels for differentially expressed genes across all clones.

**Extended Data Fig. 7.**
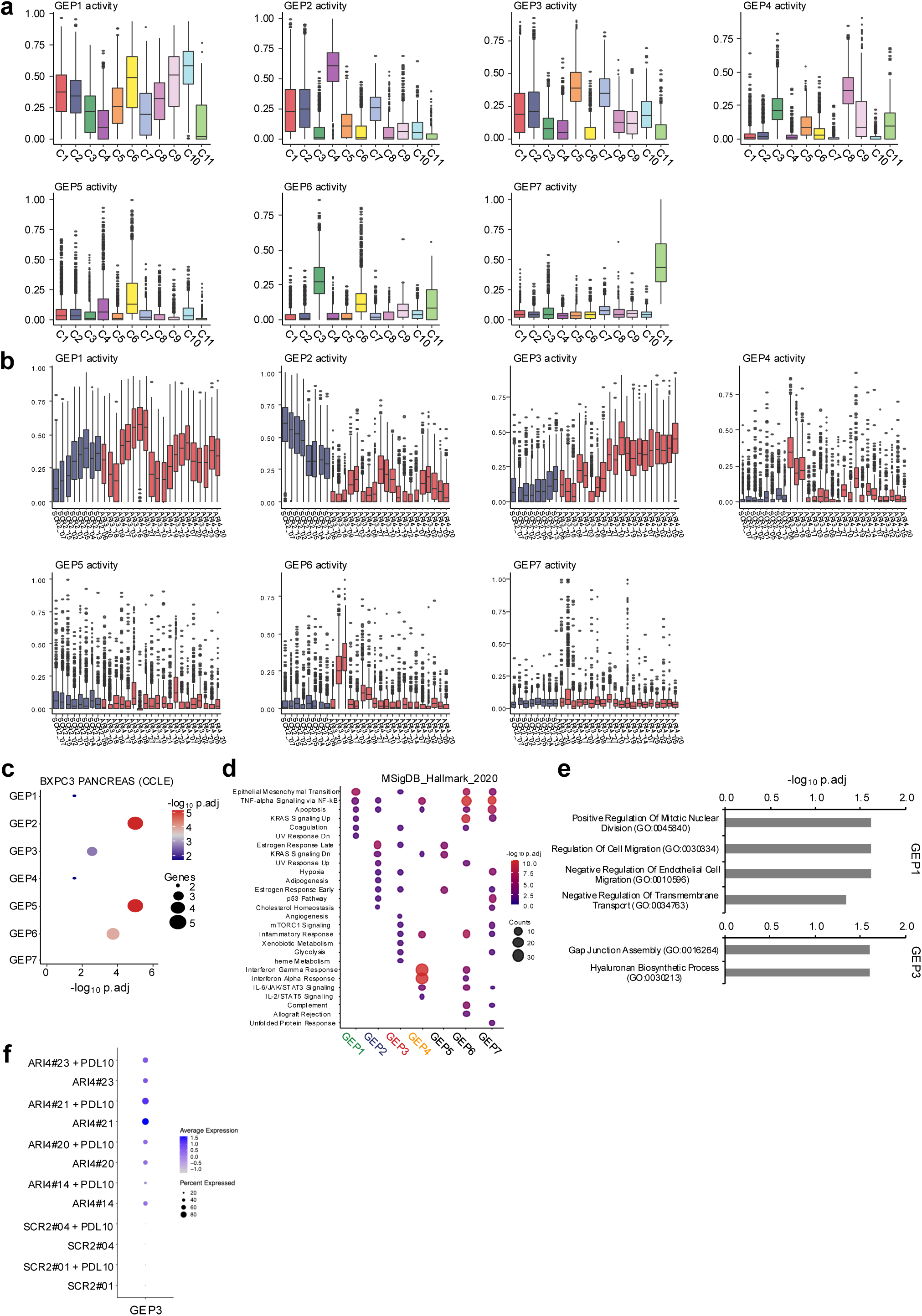
Gene expression program (GEP) activity and functional associations across 28 clones analyzed by IMPACH. (**a, b**) Box plots showing the activity scores of seven gene expression programs (GEP1–GEP7) across 28 clones, grouped by transcriptomic clusters (**a**, top) and by individual clones (**b**, bottom). (**c**) Dot plot showing the enrichment of each GEP’s constituent gene against BxPC3 marker genes listed in the Cancer Cell Line Encyclopedia (CCLE). The dot size represents the number of overlapping genes; color indicates −log₁₀ adjusted *P*-values. (**d**) Dot plot showing enrichment of MSigDB Hallmark 2020 gene sets across GEP1–GEP7. Dot size reflects the number of overlapping genes; color scale denotes −log₁₀ adjusted *P*-values. (**e**) Gene Ontology (GO) enrichment analysis (Biological Process category) for genes in GEP1 (top) and GEP3 (bottom). Bar plots show the most significantly enriched GO terms ranked by −log₁₀ adjusted *P*-values. (**f**) Dot plot showing the average expression and detection frequency of representative GEP3-associated genes across individual clones. GEP3 assignment was based on the expression profiles of approximately 10 cell divisions after the initial molecular profiling. PDL, population doubling level.

**Extended Data Fig. 8.**
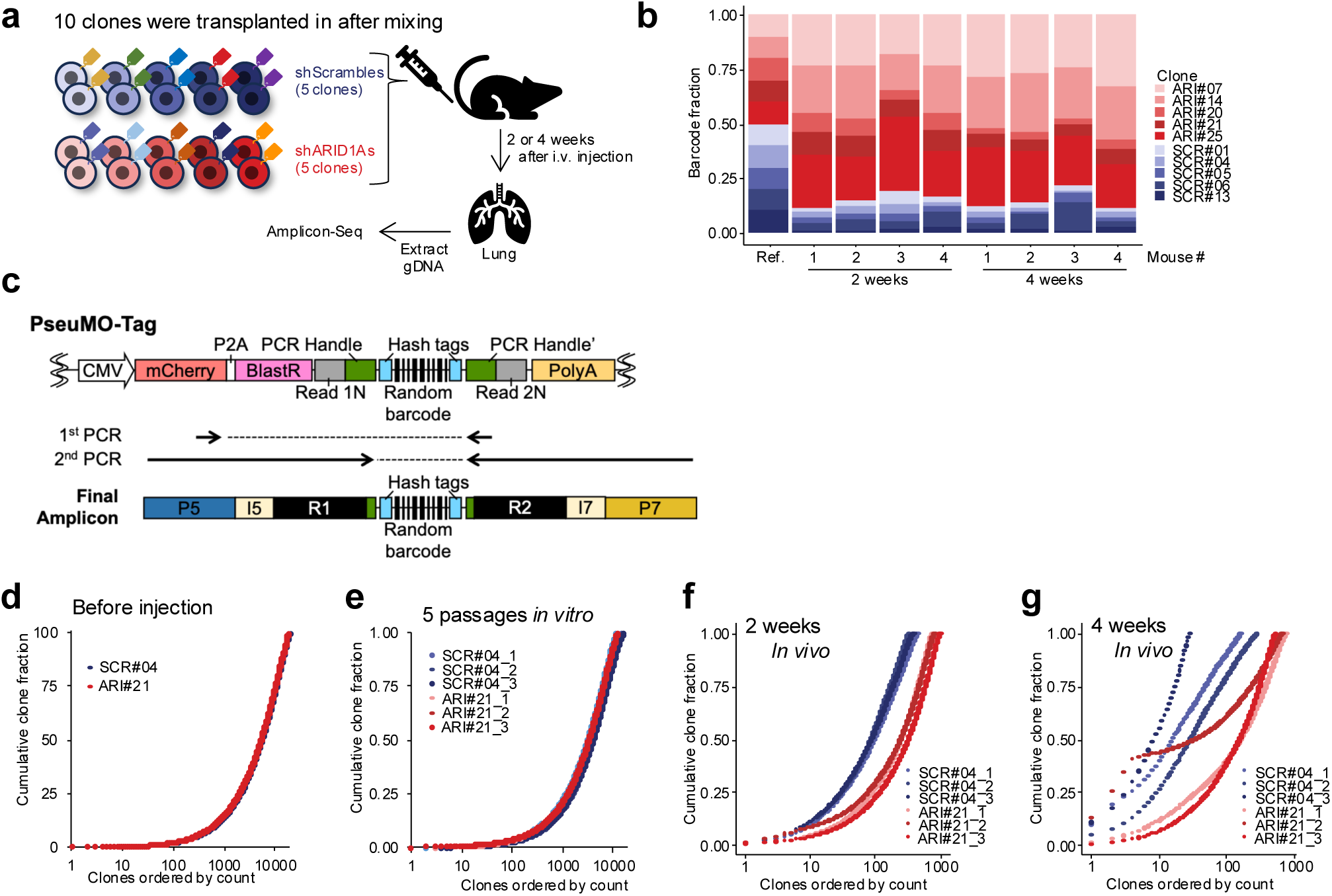
Clonal dynamics of shPseuMO-Tag–labeled BxPC3 clones *in vitro* and *in vivo*. (**a**) Schematic of the *in vivo* transplantation assay. Pooled mixture of five shScramble and five shARID1A BxPC3 clones was intravenously injected into NSG mice. Lung tissues were harvested at 2 or 4-weeks post-injection for clonal barcode analysis. (**b**) Stacked bar plots showing the relative barcode fractions of individual clones recovered from the lung tissues of recipient mice at 2 and 4 weeks. The reference (input) clone distribution is shown for comparison. (**c**) Schematic of the PseuMO-Tag vector and PCR strategy for barcode amplification. (**d**, **e**) Cumulative clone fraction plots showing clonal distributions *in vitro* before injection (d) and after five passages in culture (e). (**f**, **g**) Cumulative clone fraction plots showing *in vivo* clonal dynamics at 2 weeks (f) and 4 weeks (g) post-injection. Red curves indicate shARID1A clones. Blue curves indicate shScramble clones.

**Extended Data Fig. 9.**
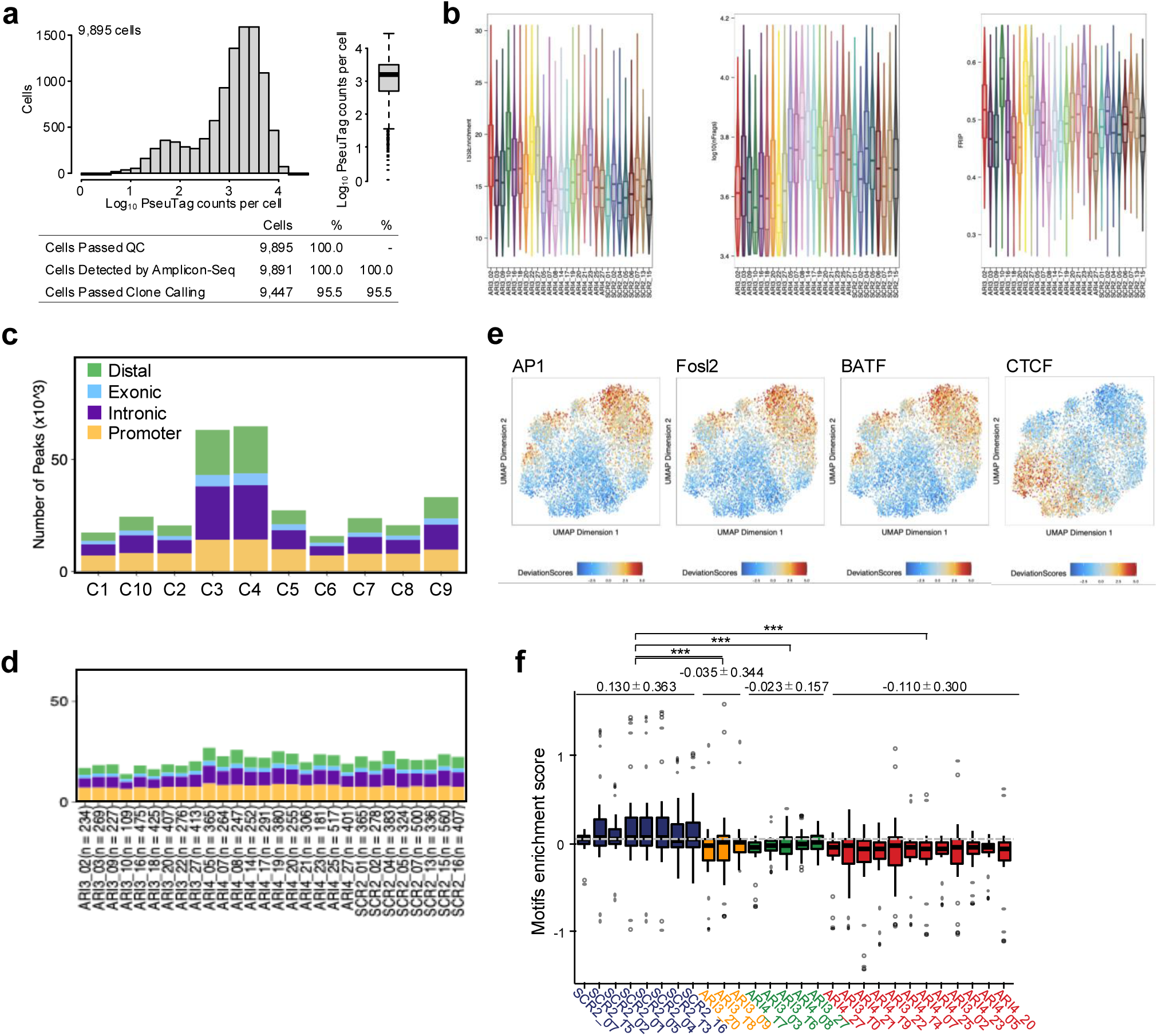
Chromatin accessibility profiling and motif analysis in 28 clones analyzed by IMPACH. (**a**) Summary of PseuMO-Tag-based clone calling from scATAC-seq data (9,985 cells). Histogram and box plot show log₁₀-transformed barcode counts per cell. The accompanying table summarizes quality control (QC) metrics, including Amplicon-seq detection rates and clone assignment rates. (**b**) Violin plots showing QC metrics from scATAC-seq data, including the number of fragments per cell (nFrags), transcription start site (TSS) enrichment scores, and fraction of reads in peaks (FRiP), stratified by clone identity. (**c**) Genomic annotation of accessible chromatin regions for each chromatin cluster, categorized as promoter, intronic, exonic, or distal elements. (**d**) Bar plots showing the number of accessible peaks per clone, annotated by genomic region. (**e**) UMAP projections colored by deviation scores of representative transcription factor motifs, calculated using chromVAR. (**f**) Motif enrichment scores across clones. Each point represents a motif. The comparisons between clones reveal statistically significant differences (\*\*\**P < 0.001*) as determined by one-way ANOVA followed by Dunnett’s post hoc multiple comparisons test. Mean ± s.d. values of enrichment scores are shown above each group.

**Extended Data Fig. 10.**
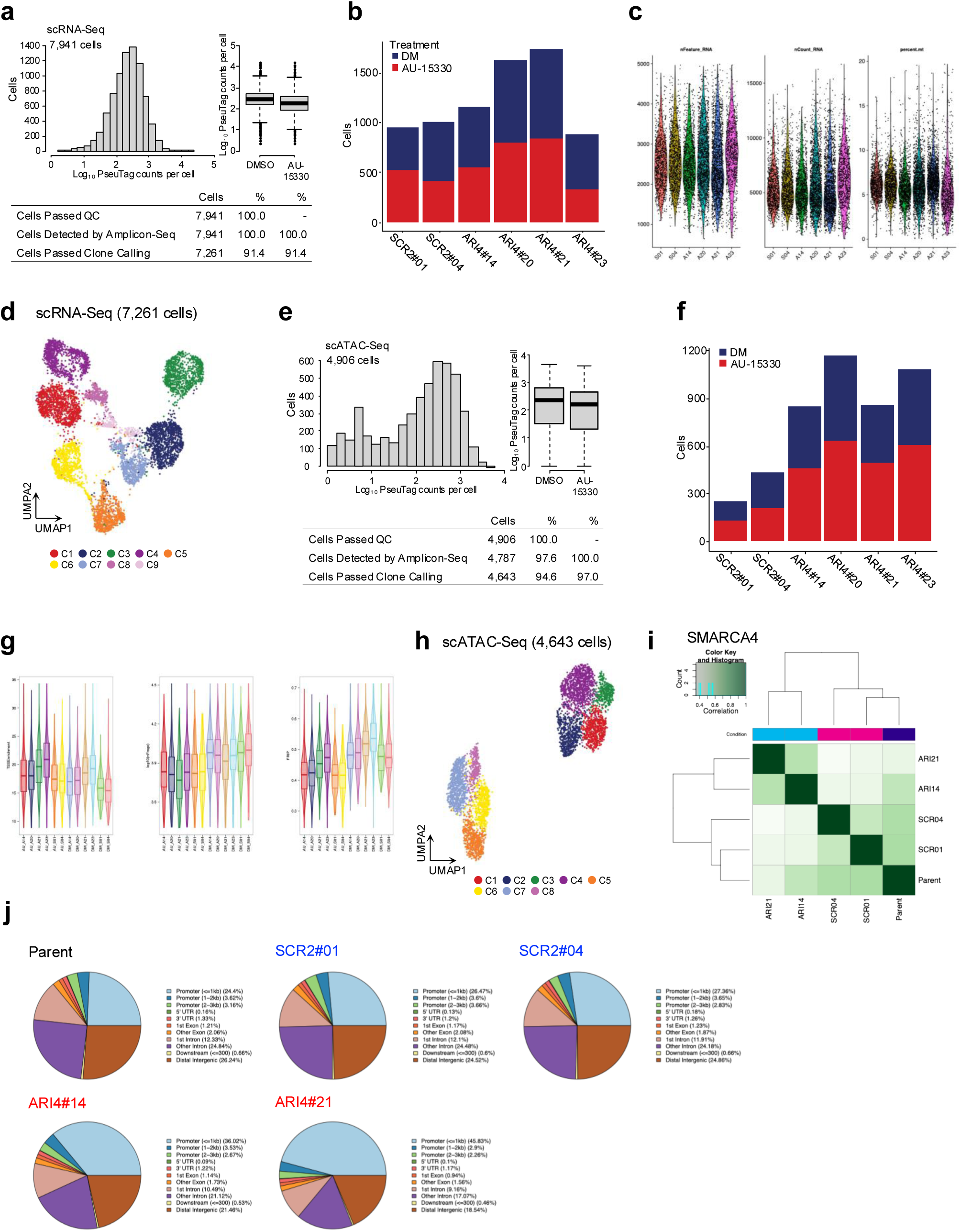
Single-cell transcriptomic and chromatin accessibility analysis of BxPC3 clones following SMARCA4 inhibition. **(a, e)** Summary of clone calling results from scRNA-seq (**a**, 7,941 cells) and scATAC-seq (**e**, 4,906 cells) datasets following treatment with either DMSO or SMARCA4 inhibitor AU-15330. Histograms and box plots showing the distribution of log₁₀-transformed PseuMO-Tag counts per cell. Tables summarize quality control (QC) metrics, including Amplicon-seq detection rates and successful clone assignment rates. **(b, f)** Bar plots showing the number of cells retained for downstream analysis under each treatment condition (DMSO or AU-15330) for scRNA-seq (**b**) and scATAC-seq (**f**). **(c, g)** Violin plots displaying standard QC metrics for scRNA-seq (**c**), including the number of detected genes (nFeature_RNA), UMI counts, and the percentage of mitochondrial RNA, and for scATAC-seq (**g**), including the number of fragments per cell (nFrags), transcription start site (TSS) enrichment scores, and fraction of reads in peaks (FRiP). **(d, h)** UMAP projections of single-cell transcriptomic (**d**) and chromatin accessibility (**h**) profiles, colored according to the unsupervised clustering results. **(i)** Heatmap depicting pairwise Pearson correlation coefficients between SMARCA4-enriched peaks across clones treated with DMSO or AU-15330. **(j)** Pie charts illustrating the genomic distribution of SMARCA4-enriched peaks (promoter, exonic, intronic, or distal regions) in parental cells and representative clones under DMSO and AU-15330 treatment conditions.

